# Identification of Multiple Iron Uptake Mechanisms in *Enterococcus faecalis* and Their Relationship to Virulence

**DOI:** 10.1101/2022.11.04.515265

**Authors:** Debra N. Brunson, Cristina Colomer-Winter, Ling Ning Lam, José A. Lemos

## Abstract

Among the unfavorable conditions bacteria encounter within the host is restricted access to essential trace metals such as iron. To overcome iron deficiency, bacteria deploy multiple strategies to scavenge iron from host tissues with abundant examples of iron acquisition systems being implicated in bacterial pathogenesis. Yet, the mechanisms utilized by the major nosocomial pathogen *Enterococcus faecalis* to maintain intracellular iron balance are poorly understood. In this report, we conducted a systematic investigation to identify and characterize the iron acquisition mechanisms of *E. faecalis* and to determine their contribution to virulence. Bioinformatic analysis and literature surveys revealed that *E. faecalis* possesses three conserved iron uptake systems. Through transcriptomics, we discovered two novel ABC-type transporters that mediate iron uptake. While inactivation of a single transporter had minimal impact on the ability of *E. faecalis* to maintain iron homeostasis, inactivation of all five systems (Δ5Fe strain) disrupted intracellular iron homeostasis and considerably impaired cell growth under iron-deficiency. Virulence of the Δ5Fe strain was generally impaired in different animal models but showed niche-specific variations in mouse models, leading us to suspect that heme can serve as an iron source to *E. faecalis* during mammalian infections. Indeed, heme supplementation restored growth of Δ5Fe under iron-depletion and virulence in an invertebrate infection model. Collectively, this study reveals that the collective contribution of five iron transporters promotes *E. faecalis* virulence and that the ability to acquire and utilize heme as an iron source is critical to the systemic dissemination of *E. faecalis*.

## INTRODUCTION

A resident of the gastrointestinal tract of animals and humans, *Enterococcus faecalis* is also a major opportunistic pathogen which includes but are not restricted to central line associated bloodstream infections (CLABSI), infective endocarditis, catheter associated urinary tract infections (CAUTI), and wound infections (1). Over the past several decades, the haphazard prescription of antibiotics combined with the intrinsic hardy nature of *E. faecalis*, including natural and acquired resistance to antibiotics, have contributed to a sustained and often times increased presence of enterococcal infection outbreaks in healthcare settings or in the community (2). Generally considered a low-grade pathogen due to the limited number of tissue damaging factors encoded in its core genome, the virulence potential of *E. faecalis* is thought to derive from a capacity to form robust biofilms on tissues or on indwelling devices, to thrive under a variety of adverse environmental conditions, and to subvert the immune system (3-5). Therefore, a better understanding of the mechanisms utilized by *E. faecalis* to survive under unfavorable conditions, especially those encountered within the human host, can potentially provide new therapeutic leads.

Among the adverse conditions pathogens encounter during infection is limited access to essential trace metals, in particular iron, manganese, and zinc that are actively sequestered by metal-binding host proteins as part of an antimicrobial process known as nutritional immunity (6-10). Iron is of particular significance as it is the preferred metal cofactor of enzymes that carry out fundamental cellular processes such that it plays a central role in host pathogen interactions (6, 11, 12). Despite being the most abundant trace metal in vertebrate tissues, iron is not readily available to bacterial pathogens because the vast majority of this element found in the host is complexed to heme inside red blood cells or bound to ferritin, an intracellular protein produced in hepatocytes that serves as the principal iron storage protein in mammalian cells (13). In addition, several host-produced proteins avidly bind free iron either to avoid iron toxicity to host tissues or as part of the nutritional immunity process (12, 14, 15). For instance, the liver produces and secretes transferrin (TF), which binds free Fe^+3^ in the bloodstream and at sites of infection, which is then recycled by macrophages by unloading iron to intracellular ferritin and returning apo-TF into circulation (13). In mucosal surfaces, free iron is sequestered by lactoferrin that is also found in high concentrations in human secretions such as saliva (16). While primarily known for its role in manganese and zinc sequestration, neutrophil-secreted calprotectin has been shown to efficiently chelate Fe^2+^ in anaerobic environments *in vivo* (9, 17). All these factors combined with the low solubility of Fe^+3^ in sera make free iron concentrations within vertebrates to be several orders of magnitude below the concentration range required for microbial growth (12, 18, 19).

To overcome host-imposed iron starvation, bacterial pathogens deploy multiple strategies to scavenge free iron directly or intracellularly stored, bound to organic molecules (such as heme) within hemoproteins, or mobilized to iron-binding proteins (6, 10, 20-22). Perhaps the most effective strategy utilized by bacteria to scavenge iron is via the production of siderophores (“iron carrier” from the Greek), which are low molecular mass organic molecules that are among the strongest metal chelators known to date (23, 24). While not all bacteria synthesize siderophores, high affinity surface-associated iron transporters are ubiquitous in bacteria with some of the most successful blood borne pathogens encoding at least one dedicated heme acquisition system in addition to elemental iron transporters (19, 21, 25, 26). Not surprisingly, many of the genes associated with siderophore biosynthesis and uptake as well as iron and heme transporters have been directly implicated in bacterial virulence (6, 25, 27-30). In recent years, our group identified and characterized the manganese and zinc import systems of *E. faecalis* showing that the well-coordinated activity of either manganese (EfaCBA, MntH1 and MntH2) or zinc (AdcABC and AdcAII) transporters is critical to *E. faecalis* fitness and virulence (31, 32). However, when it comes to the mechanisms utilized by the enterococci to maintain iron homeostasis and its relationship to enterococcal pathogenesis, current knowledge is restricted to *in silico* and transcriptome-based studies showing that *E. faecalis* encodes three highly conserved iron import systems that are regulated by either the DtxR-like/EfaR repressor (EfaCBA) or the Fur-like repressor (FeoAB and FhuDCBG) (33-35). To fill this current knowledge gap, we sought in this study to identify and characterize the mechanisms utilized by *E. faecalis* to overcome iron starvation and determine the individual and collective contributions of iron uptake systems to *E. faecalis* virulence. Through transcriptomics, we identified two additional and previously uncharacterized ABC-type iron transporters restricted to enterococci and a limited number of streptococcal species. We named the novel iron transporters FitABCD and EmtABC and generated strains lacking one or both transporters using the Δ*fitABΔemtB* double mutant as the background to generate a quintuple mutant also lacking *efaCBA, feoAB* and *fhuDCBG* (Δ5Fe strain). Characterization of these mutant strains revealed that *E. faecalis* indeed utilizes multiple iron transporters to acquire iron under iron-depleted conditions and that their collective activity is important for enterococcal pathogenesis in a niche-specific manner. In addition, evidence that *E. faecalis* can utilize heme as an alternative iron source and that unidentified heme transporter(s) might be critical for systemic dissemination and disease outcome is also provided.

## RESULTS

### Two uncharacterized ABC-type transporters are the most upregulated genes in *E. faecalis* OG1RF grown under iron-depleted conditions

To identify the genes and pathways utilized by *E. faecalis* to grow under iron starvation, we used RNA deep sequencing (RNA-seq) to compare the transcriptome of the parent strain OG1RF grown to mid-log in the chemically defined FMC medium (31, 36) with or without the addition of FeSO_4_ as an iron source (Table S1). Despite the ∼1600-fold difference in iron content of the two media formulations (∼80 μM total iron in FMC[+Fe] compared to ∼0.05 μM total iron in FMC[-Fe], Table 1), the ability of different *E. faecalis* and *E. faecium* strains to grow under iron-replete or iron-depleted conditions was remarkably similar (Fig. 1). Moreover, quantification of intracellular elemental iron in the *E. faecalis* OG1RF strain grown to mid-log phase in FMC[+Fe] or FMC[-Fe] revealed a small and not statistically significant difference between the two conditions (0.410±0.122 μM intracellular iron in FMC[+Fe] versus 0.322±0.127 μM iron in FMC[-Fe]). These results strongly indicate that the enterococci are well equipped to scavenge iron and maintain iron homeostasis under extreme conditions. To facilitate interpretation of the RNA-seq study, we used a false discover rate (FDR) of 0.01 and applied a 2-fold cutoff to generate a list of differently expressed genes (Table S2). For illustration purposes, the 200 differentially expressed genes (92 upregulated and 108 downregulated) were grouped according to Clusters of Orthologous Groups (COG) functional categories, with genes coding for membrane-associated transporters (22%), metabolism (31%), and hypothetical proteins (53%) comprising the majority of genes identified in the comparison (Fig. 2). When compared to cells grown in FMC[+Fe], the most upregulated genes (varying from 2.6- to 7.7-fold induction) in cells grown in FMC[-Fe] coded for proteins that belong two uncharacterized ABC-type transport operons (OG1RF_RS12045 to OG1RF_12060 and OG1RF_RS12585 to OG1RF_12595) (Table S2, Fig. 3). While there is no previous experimental evidence that these transporters are involved in metal uptake, OG1RF_RS12045-12060 was previously shown to be part of the Fur (ferric <u>u</u>ptake regulator) regulon (33) and is presently annotated as putative ABC-type iron transporter (35). Herein, we will refer to OG1RF_RS12045-12060 as *fitABCD* for <u>F</u>ur regulated iron transporter and OG1RF_RS12585-12595 as *emtABC* for <u>e</u>nterococcal <u>m</u>etal transporter. Based on searches of public databases and phylogenetic tree analyses with the substrate binding proteins FitD or EmtC, the proteins encoded by the *fitABCD* and *emtABC* are highly conserved among the enterococci (Fig. 3 and Fig. 4). Beyond enterococci, FitD shares ∼ 48% amino acid identity and ∼ 65% similarity with the *B. subtilis* YclQ and *S. pneumoniae* SPD_RS08810 whereas the non-enterococcal protein most closely related to EmtC is the *S. pyogenes* RS01525 that shares 29% identity and 47% similarity with EmtC. Notably, the *Bacillus subtilis* YclNOPQ has been implicated in the uptake of the petrobactin siderophore (33, 37) such that it is possible that EitABCD is involved in the uptake of siderophore. Other than the upregulation of *fitABCD* and *emtABC* operons, few other notable alterations in the iron starvation transcriptome were the upregulation of genes from the mannose PTS and pyrimidine biosynthesis operons and the downregulation of two P-type ATPases annotated as magnesium import transporters and the tellurite (toxic anion) resistance protein (Table S2). While studies to understand the significance of these other notable transcriptional changes to growth under iron starvation were not pursued in this study, these changes are suggestive of adaptation to iron starvation triggering changes in carbon and nucleic acid metabolism and metal resistance profiles.

**TABLE 1.** Iron and manganese quantifications in the media used in this study.

| Media | Fe (μM) | Mn (μM) |
| --- | --- | --- |
| FMC | 80.02 (+/- 10.123) | 118.401 (+/-16.265) |
| FMC [-Fe] | 0.052 (+/-0.008) | 109.881 (+/-6.870) |
| FMC Low-Metal (FMC-LM) | 10.538 (+/- 0.829) | 10.414 (+/-0.807) |
| FMC-LM [-Fe] | 0.002 (+/-0.0048) | 14.853 (+/-3.362) |
| FMC [-Fe/-Mn] | 0.0376(+/-0.103) | 0.0413 (+/-0.0215) |
| FMC [-Fe/+heme] | 4.584 (+/-0.182) | 11.992 (+/-3.811) |
| BHI | 6.581 (+/- 0.950) | 0.570 (+/-0.0208) |

**FIG 1.**
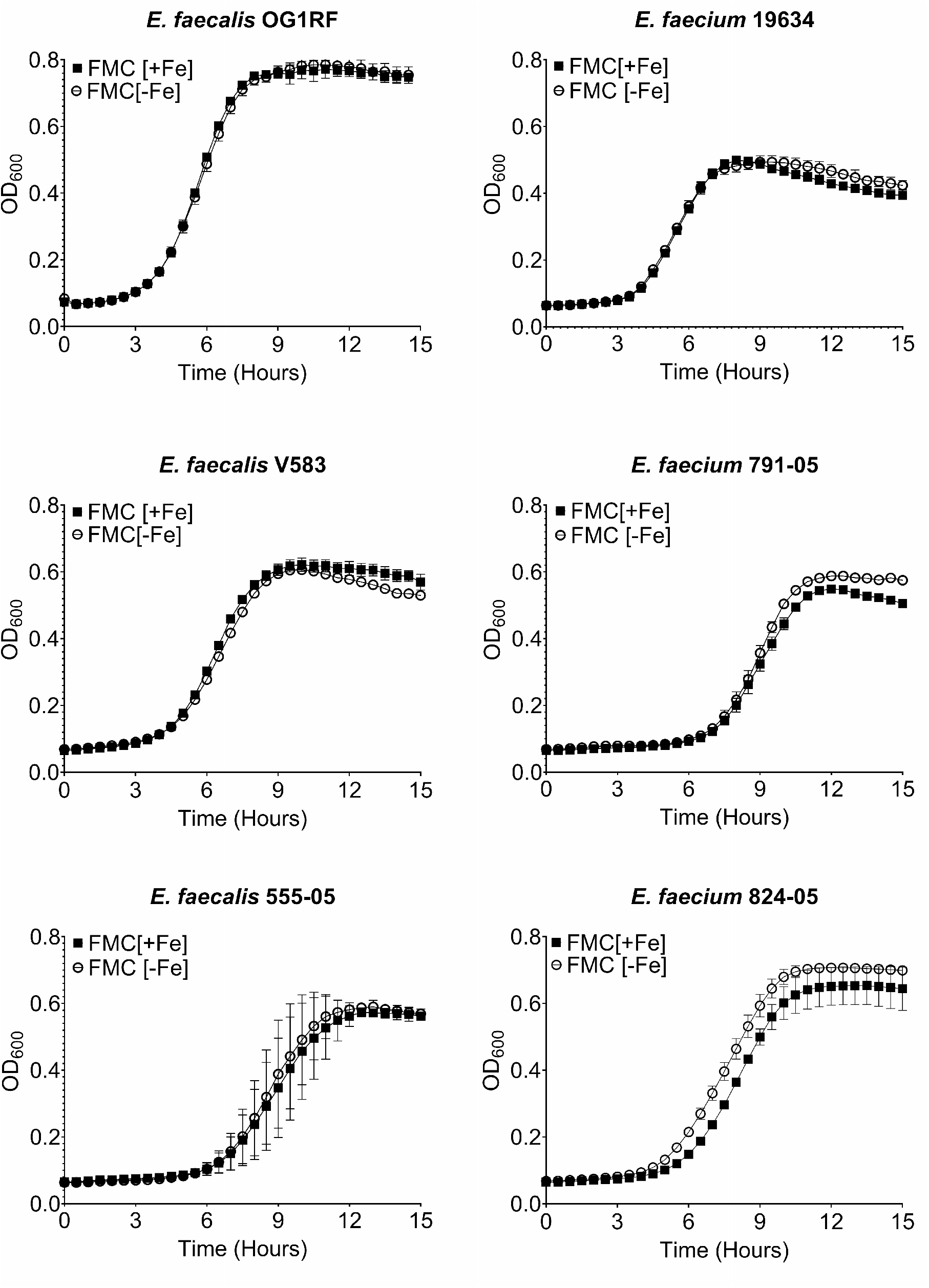
Growth of *E. faecalis* and *E. faecium* strains in FMC[+Fe] or FMC[-Fe]. Growth was monitored by measuring OD_600_ every 30 minutes using an automated growth reader. Error bars denote standard deviations from three independent biological replicates.

**FIG 2.**
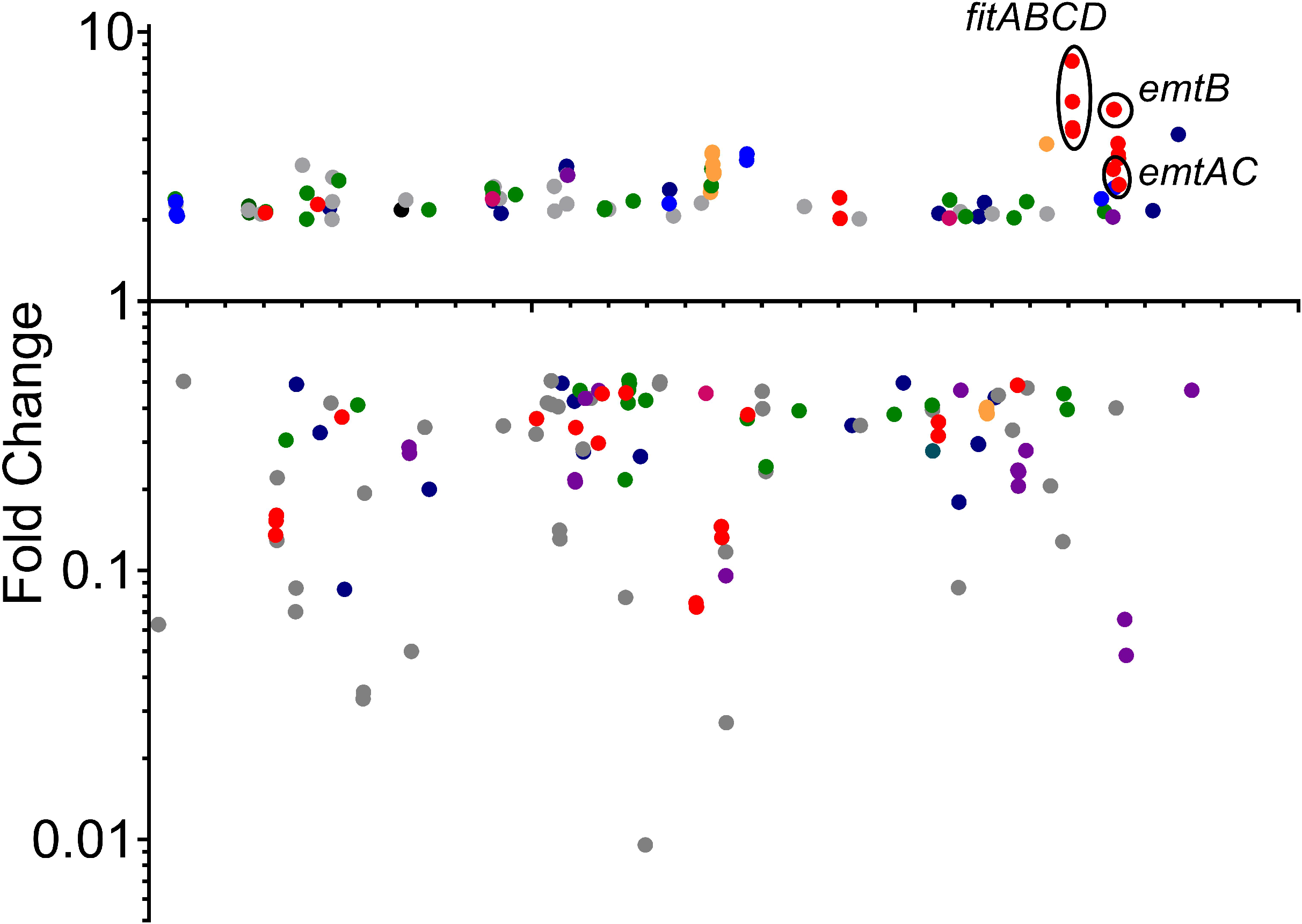
Summary of RNA-Seq analysis comparing *E. faecalis* grown under iron depleted versus iron replete conditions. Dot plot of genes differently expressed, via RNA sequencing, under conditions of iron depletion as determined by Degust (degust.erc.monash.edu). The *y* axis indicates the fold change in expression compared to control cultures, while the *x* axis indicates position within the genome.

**FIG 3.**
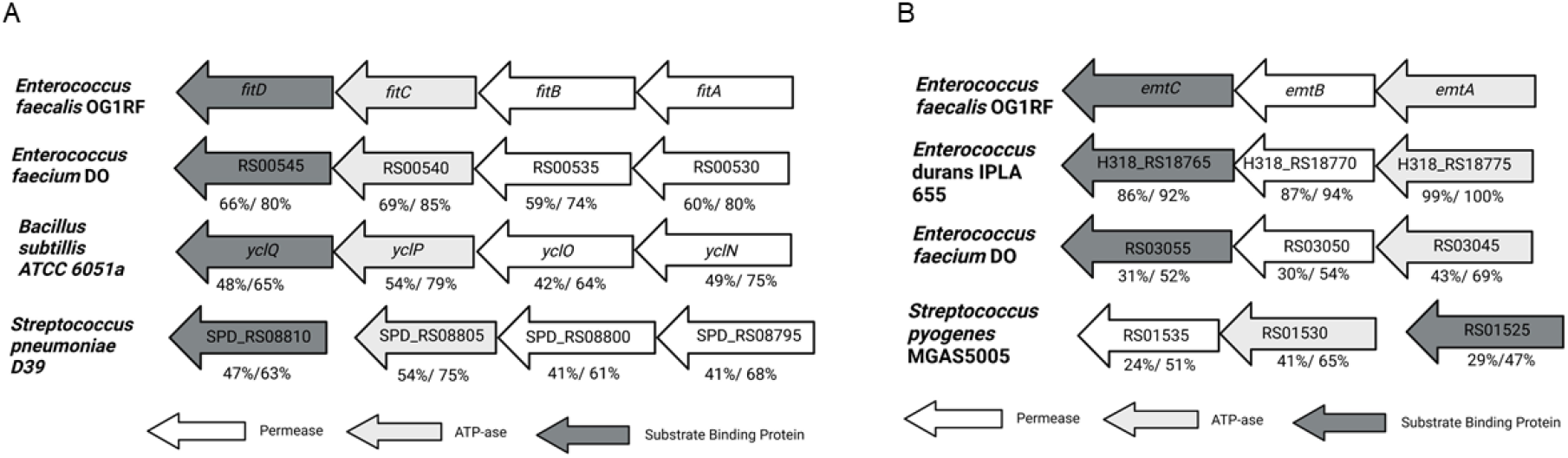
Genetic organization of FitABCD (A) and EmtABC (B) and homologues found in selected Gram-positive bacteria. Percentage of amino acid identity and positive identity to OG1RF for substrate binding protein, permease, and ATPase are indicated.

**FIG 4.**
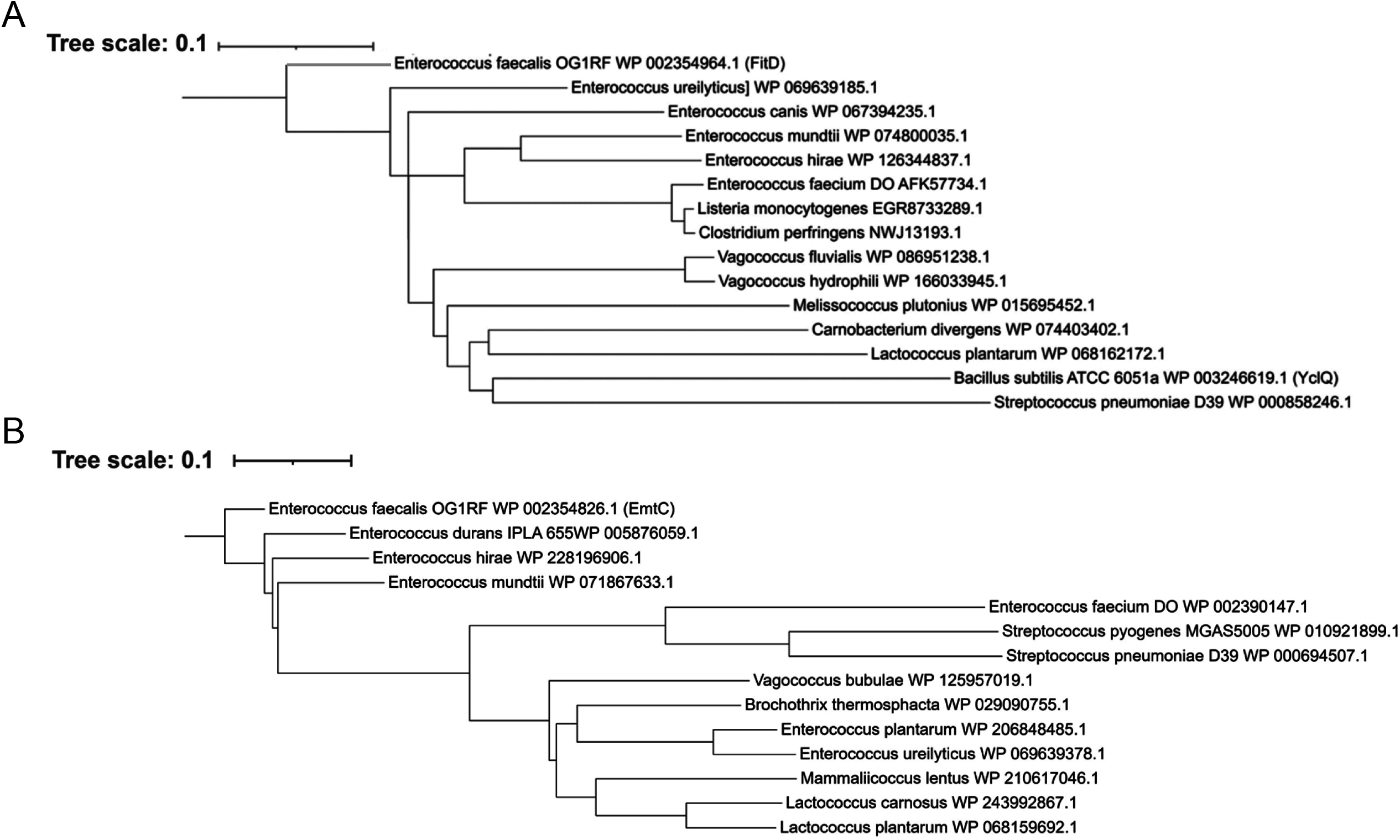
Phylogenetic tree analysis of the substrate binding proteins FitD (A) and EmtC (B). BLASTP searches against FitD and EmtC were used to identify homologues across species of enterococci, streptococci, bacilli, and other Gram-positive bacteria. Phylogenetic trees were constructed using multiple sequence alignments of representative species using Clustal Omega and iTOL.

### FitABCD and EmtABC are important but not critical for growth under low iron conditions

To determine the contributions of FitABCD and EmtABC to growth of *E. faecalis* under replete or depleted iron conditions, each system was inactivated alone or in combination and the ability of Δ*fitAB*, Δ*emtB* and Δ*fitAB*Δ*emtB* strains to grow in media containing different concentrations of iron and manganese assessed. In Brain Heart Infusion (BHI), a complex media with ∼ 6.5 μM iron, all mutants grew as well as the parent strain OG1RF (Fig. 5A). In (chemically-defined) FMC, which contains high concentrations of iron (75 μM FeSO_4_) and manganese (100 μM MnSO_4_) in the original recipe (36), all strains grew well although Δ*fitAB* and Δ*fitAB*Δ*emtB* attained slightly lower final growth yields (Fig. 5B). The omission of FeSO_4_ from FMC slightly delayed growth and further lower final growth yields of Δ*fitAB* and Δ*fitAB*Δ*emtB* as well as Δ*emtB* (Fig. 5C). Because iron and manganese may function as interchangeable cofactors and *E. faecalis* is deemed a “manganese-centric” organism (31), we prepared a modified low metal FMC (LM-FMC) formulation containing 1/10^th^ of the original concentrations of iron and manganese for subsequent studies (Table 1). Like the original FMC recipe, the Δ*fitAB* and Δ*fitAB*Δ*emtB* strains reached lower final growth yields in complete LM-FMC with all mutants growing more poorly in LM-FMC[-Fe] (Fig. 5D-E). Finally, all strains (parent strain included) grew slower and reached lower final growth yields in LM-FMC lacking both iron and manganese (Fig. 5F). Collectively, these results indicate that FitABCD and EmtABC contribute but are not essential to growth under iron-depleted conditions.

**FIG 5.**
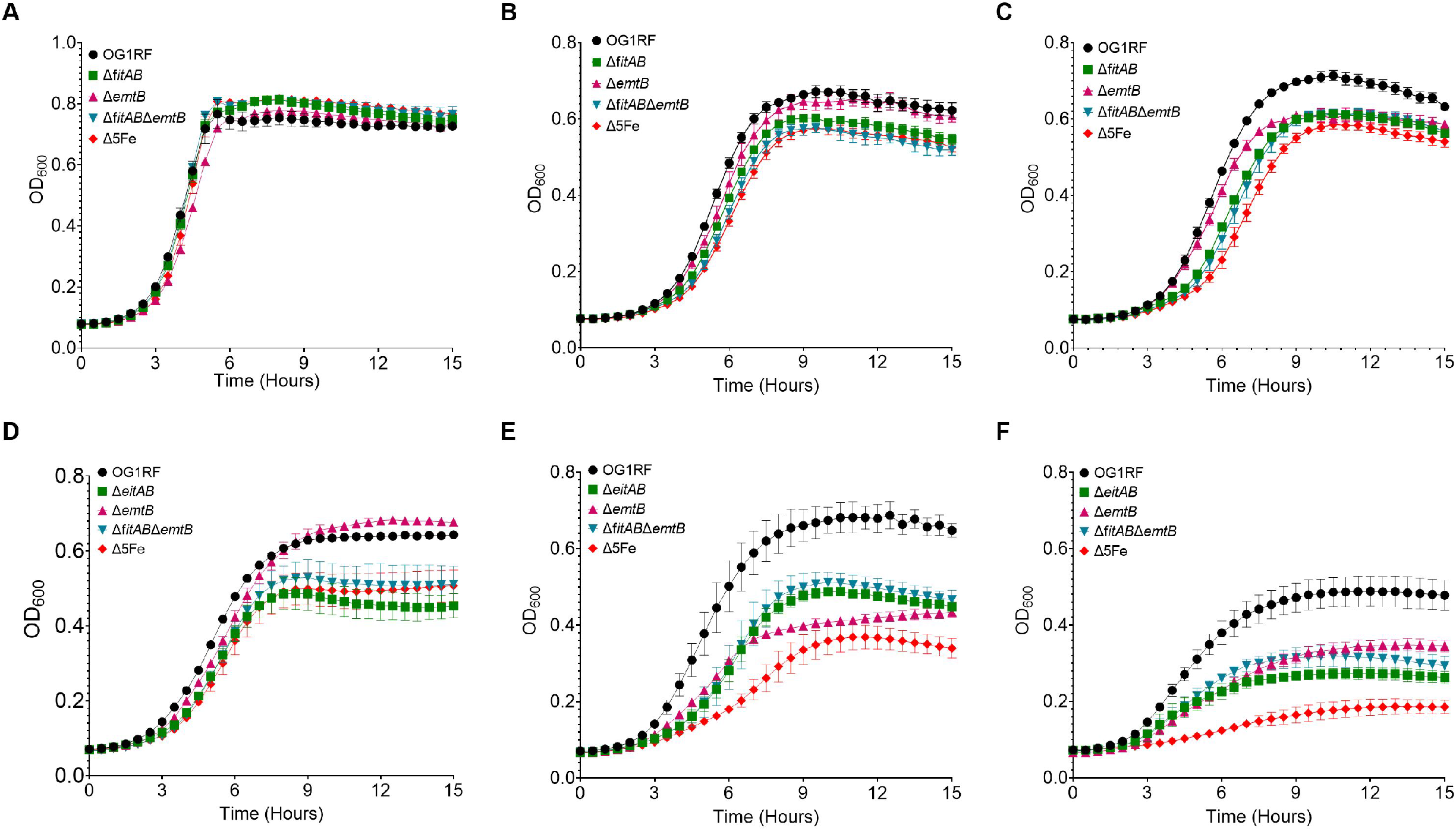
Growth of OG1RF, Δ*fitAB*, Δ*emtB*, Δ*fitAB*Δ*emtB*, and Δ5Fe in (A) BHI, (B) FMC[+Fe], (C) FMC[-Fe], (D) LM-FMC[+Fe], (E) LM-FMC[-Fe], and (F) LM-FMC[-Fe/-Mn]. Growth was monitored by measuring OD_600_ every 30 minutes using an automated growth reader. Error bars denote standard deviations from three biological replicates.

### Temporal expression of iron transporters in response to iron starvation

Previous studies revealed that the conserved iron transporters *feoAB* and *fhuDCBG* are regulated by the iron-sensing Fur regulator (33) whereas transcription of the dual iron/manganese transporter *efaCBA* is controlled by the manganese-sensing EfaR regulator (38). While none of the genes from the *feoAB, fhuDCBG* and *efaCBA* operons were differently expressed in our RNA-seq analysis, we suspected that their transcriptional activation in response to iron starvation may occur immediately after cells are starved for iron returning to basal expression levels after cells have become adapted to the new (low iron) environment. To investigate this possibility, we monitored (via reverse transcriptase quantitative PCR, RT-qPCR) the transcriptional pattern of *efaCBA, feoAB, fhuDCBG* as well as *fitABC* and *emtABC* within the first hour after cultures were switched from iron replete to iron depleted condition. Using one representative gene for each operon as proxy, we found that all transcriptional units were upregulated in response to iron starvation (Fig. 6). Noteworthy, this induction occurred in two distinctly separated surges. In the first surge appeared *emtB* and *efaA* that were strongly induced 10-min after cells were starved for iron but returning to near basal levels of expression after 60-min. In the second surge, *fitA* and *fhuB* were much more strongly induced at the later (60-min) time point. Finally, transcription of *feoB* was not altered during the initial 30 minutes but displayed a modest (yet significant) upregulation at 60-min such that considered *feoAB* part of the second surge. These results strongly suggest that *E. faecalis* encodes, at the minimum, five bona-fide iron import systems that can be grouped into early and late responders.

**FIG 6.**
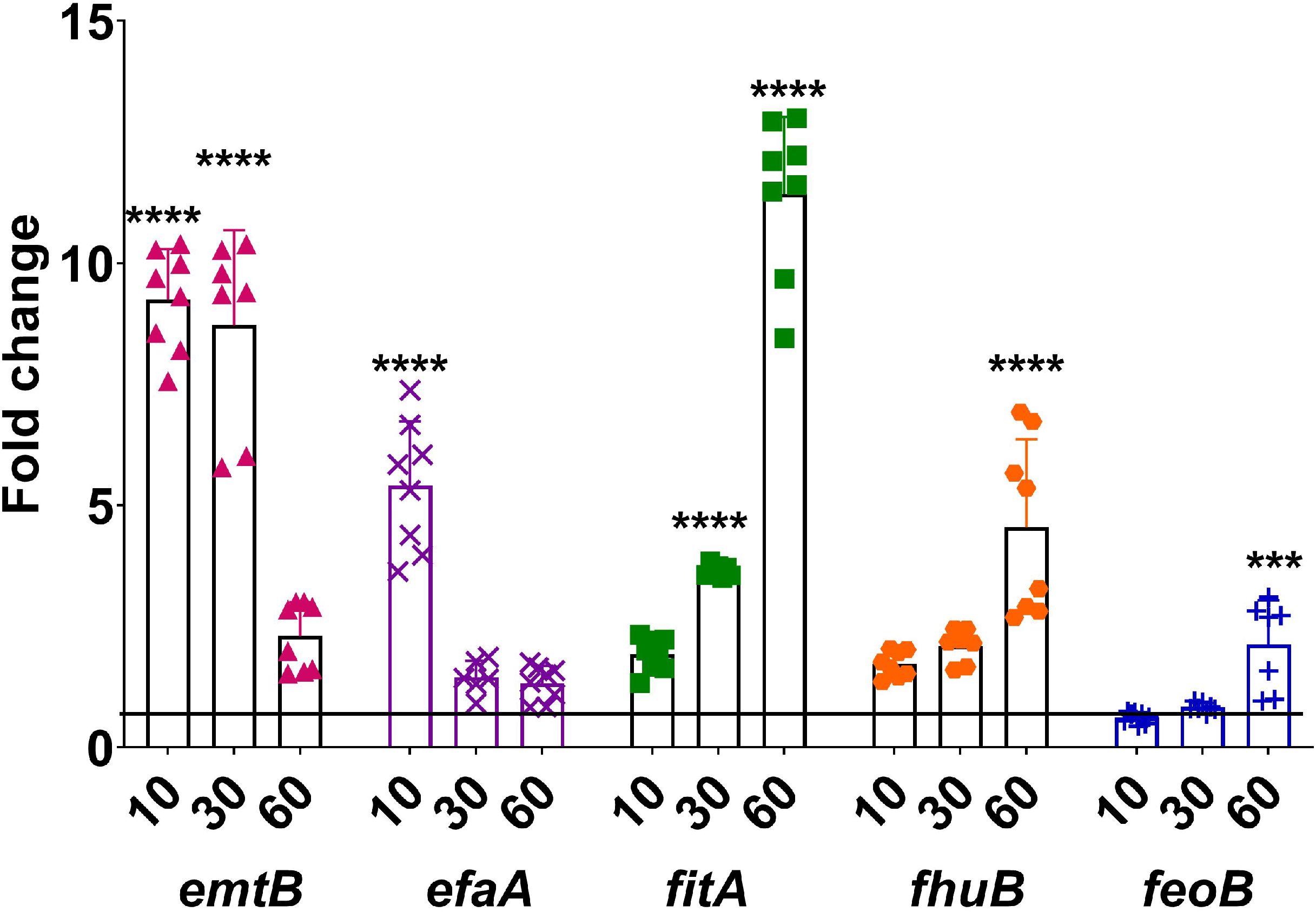
Quantitative real time PCR analysis of iron transport genes transferred from iron replete (FMC[+Fe]) to iron depleted (FMC[-Fe]) conditions. Data shown represents four independent cultures with two technical replicates each. Line is set at 1 to indicate expression equivalent at T_0_. Significance was determined by one-way ANOVA using a Dunnet’s post-hoc test to compare mRNA levels at T_0_ with T_10_, T_30_ and T_60_ time points. ***p≤0.001, and ****p≤0.0001.

### Simultaneous inactivation of *efaCBA, feoAB, fhuDCBG, fitABCD* and *emtABC* further impairs growth in iron-depleted conditions

To probe the individual and collective contributions of EfaCBA, FeoAB, and FhuDCBG to iron homeostasis, we took advantage of the Δ*efaCBA* strain that was already available in the lab (31) and isolated two new deletion mutants lacking FeoAB (Δ*feoB*) and FhuDCBG (Δ*fhuB*). In BHI, LM-FMC, LM-FMC[-Fe] or LM-FMC[-Fe and -Mn], the Δ*feoB* and Δ*fhuB* single mutants phenocopied the parent strain (Fig. S1). The Δ*efaCBA* strain also phenocopied growth of the parent strain in BHI, LM-FMC or LM-FMC[-Fe], but could barely grow in LM-FMC[-Fe and -Mn] (Fig. S1), a phenotype that can be attributed to the role of EfaCBA in the uptake of both iron and manganese (31). Because the dual role of EfaCBA in iron and manganese acquisition creates a confounding factor (impaired manganese uptake), we next isolated a Δ*feoB*Δ*fhuB*Δ*fitAB*Δ*emtB* strain by sequentially inactivating *feoB* and *fhuB* in the Δ*fitAB*Δ*emtB* background such that a functional EfaCBA is retained in this mutant. However, this quadruple mutant grew exactly like the double mutant Δ*fitAB*Δ*emtB* in either LM-FMC or LM-FMC[-Fe] (Fig. S2). For this reason, our next step was to introduce the *efaCBA* deletion in the quadruple mutant background yielding the Δ*efaCBA*Δ*feoB*Δ*fhuB*Δ*fitAB*Δ*emtB* strain, which we will call Δ5Fe strain onwards. In BHI, FMC[+/-Fe] and LM-FMC, growth of the Δ5Fe strain was comparable to the growth rates and yields obtained for all singles, double (Δ*fitAB*Δ*emtB*) and quadruple (Δ*feoB*Δ*fhuB*Δ*fitAB*Δ*emtB*) mutants (Fig. 5A-D, Fig. S1 and Fig. S2). However, the Δ5Fe strain grew slower and had lower growth yields when compared to the Δ*fitAB*, Δ*emtB*, and Δ*fitAB*Δ*emtB* strains grown in LM-FMC[-Fe] and LM-FMC[-Fe and -Mn] (Fig. 5E-F).

To further understand the specific contributions of FitABCD and EmtABC and the collective contribution of the five transporters to iron homeostasis, we used inductively coupled plasma optical-emission spectrometry (ICP-OES) to determine the intracellular iron concentrations in the parent, Δ*fitAB*, Δ*emtB*, Δ*fitAB*Δ*emtB* and Δ5Fe strains grown to mid-log phase in either LM-FMC or LM-FMC[-Fe]. In agreement with results showing that all strains grow well in iron replete media (Fig. 5), no significant differences in intracellular iron content were observed between parent and mutant strains when grown in LM-FMC (Fig. 7A). On the other hand, intracellular iron pools were significantly lower in the Δ*emtB* (*p*≤ 0.05) and Δ5Fe strains (*p*≤ 0.001) when grown in LM-FMC[-Fe]. While the ∼ 45% reduction in iron pools in the Δ*emtB* strain is apparently at odds with the results obtained with the Δ*fitAB* or double mutant strains, the ∼ 90% reduction observed for the quintuple mutant bodes well with the marked growth defect of this strain in LM-FMC[-Fe]. To complement these observations, we determined iron (^55^Fe) uptake kinetics in cultures of the parent, Δ*fitAB*Δ*emtB* and Δ5Fe strains grown to mid-log phase in LM-FMC[-Fe]. Time course monitoring of ^55^Fe uptake revealed a linear increase in iron uptake for the parent and Δ*fitAB*Δ*emtB* strains, while Δ5Fe displayed a non-linear and significantly (*p*≤ 0.01) reduced capacity to take up ^55^Fe over time (Fig. 7B).

**FIG 7.**
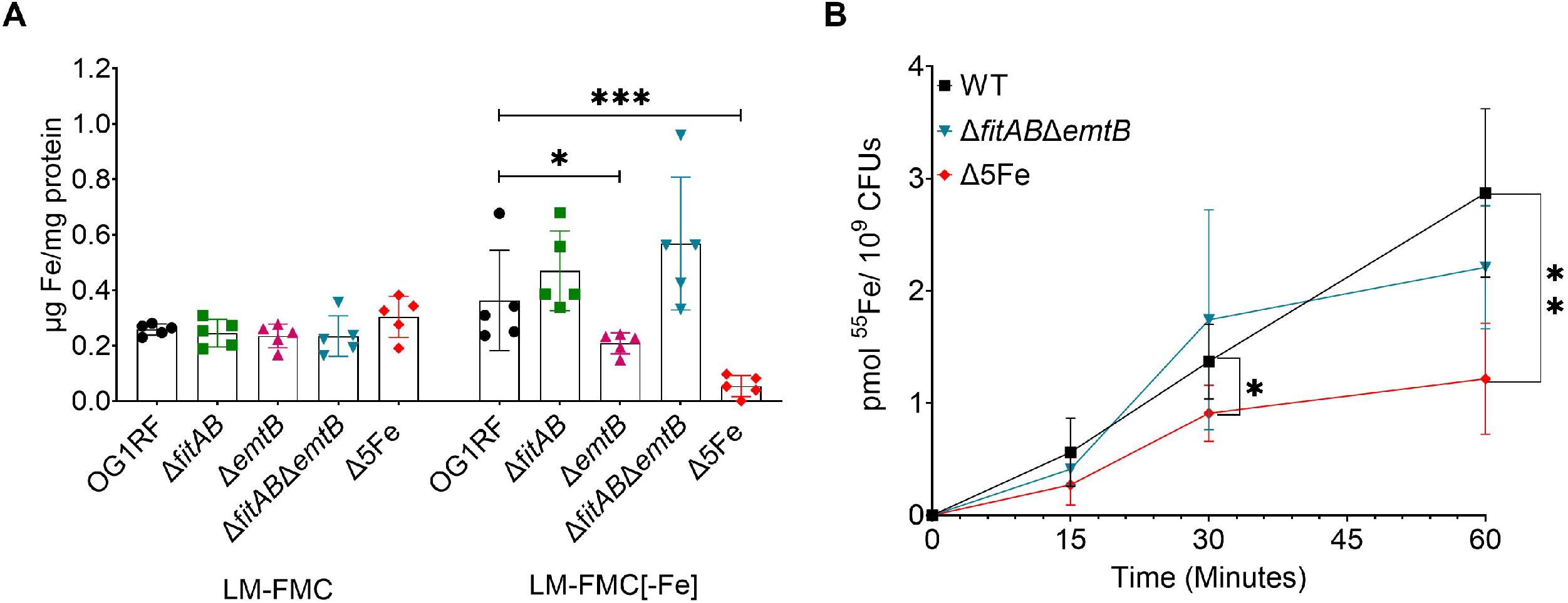
The Δ5Fe strain displays a major defect in iron acquisition. (A) ICP-OES analysis of intracellular iron content in OG1RF, Δ*fitAB*, Δ*emtB*, Δ*fitAB*Δ*emtB*, and Δ5Fe strains grown in iron replete (LM-FMC[+Fe]) or depleted (LM-FMC[-Fe]) media. (B) ^55^Fe uptake kinetics of OG1RF, Δ*fitAB*Δ*emtB* and Δ5Fe strains. Cells were grown in LM-FMC[-Fe] to mid-log phase and iron uptake monitored over time after addition of 10μM ^55^Fe. The results shown represent the average and standard deviation of five biological replicates for each data point. Significance was determined by two-way ANOVA followed by a Dunnett’s post comparison test. *p≤0.05, **p≤0.01, and ***p≤0.001.

Next, we asked if loss of FitABCD, EmtABC, or all five iron transporters affected the pathogenic potential of *E. faecalis* by testing the ability of the Δ*fitAB*, Δ*emtB*, Δ*fitAB*Δ*emtB* and Δ5Fe strains to grow and remain viable in human sera *ex vivo* as well as their virulence potential in the *Galleria mellonella* invertebrate model and in two mouse infection models. We found that, in comparison with the parent strain, the Δ5Fe strain but not Δ*fitAB*, Δ*emtB* or Δ*fitAB*Δ*emtB* was recovered in significant lower numbers after 24 hours incubation in pooled human sera at 37°C (Fig. S3). We expanded the sera growth/survival analysis by comparing the ability of parent and Δ5Fe strains to grow and then remain viable in sera for up to 48 hours. Similar to previous studies showing that mutants with defects in manganese or zinc uptake grow poorly in sera (31, 32), the Δ5Fe displayed a marked and significant growth defect in sera growing less than 1-log during the initial 12 hours of incubation compared to the parent strain that grew nearly 2-logs over the same period of time (Fig. 8A).

**FIG 8.**
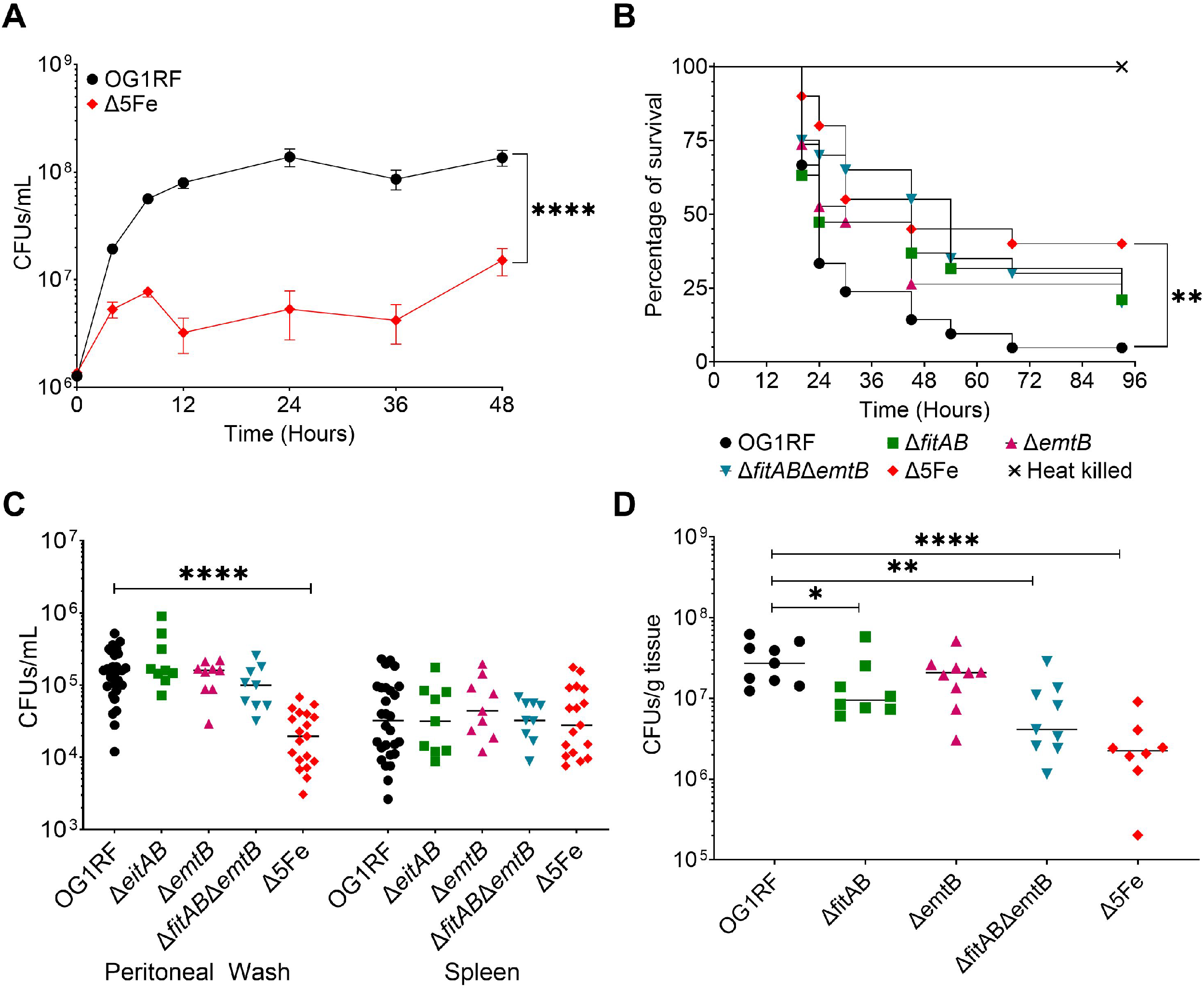
The Δ5Fe strain displays defective growth/survival in human serum *ex vivo* and attenuated virulence in animal infection models. (A) Serum was obtained from blood pooled from 3 healthy donors, bacteria were inoculated into serum at 1.5 × 10^6^ CFU, incubated at 37°C and growth/survival monitored for 48 hours. The experiment was repeated on four independent occasions with three bacterial biological replicates on each occasion. Error bars denote SEM and significance was determined using the Mann-Whitney U test. (B) Percent survival of *Galleria mellonella* infected with OG1RF, Δ*fitAB*, Δ*emtB*, Δ*fitAB*Δ*emtB*, Δ5Fe, or heat killed OG1RF. Twenty larvae were infected with 5 × 10^5^ CFU of designated strains and incubated in the dark at 37°C to monitor survival over time. The Kaplan Meyer plot is a representative of three independent experiments. Significance was determined using the Mantel-Cox log-rank test. (C) Seven-week-old C57Bl6J mice were infected via intraperitoneal injection with 2 × 10^8^ CFU of the designated strain. At 48 hours post infection, mice were euthanized, and peritoneal washes and spleens collected for CFU determination. Mann-Whitney U test was used to determine significance. (D) Seven-week-old C57Bl6 mice were wounded with a 6 mm biopsy punch and infected with 2 x10^8^ CFU of designated strains. At 3-days post infection, mice were euthanized, wounds extracted and homogenized for CFU determination. (C-D) Data points shown are a result of the ROUT outlier test and bars denote median values. Statistical analyses were performed using the Mann-Whitney test. *p≤0.05, **p≤0.01, and ****p≤0.0001.

Because trace metal sequestration is an evolutionarily conserved defense mechanism present in both vertebrates and invertebrates (39-41), previous studies conducted by our group revealed that virulence of manganese or zinc transport mutants in *Galleria mellonella* was severely compromised (31, 32), we assessed the ability of these mutants to kill *G. mellonella*. While the trends of the Kaplan-Meyer curves shown in Figure 8B are indicatives that virulence may be compromised in all the mutants tested, statistical significance (*p*≤0.01) were only achieved when comparing parent and Δ5Fe strains.

Our next step was to expand the *in vivo* studies to two mouse infections models; a peritonitis model where infection becomes systemic within 12 to 24 hours (42-44) and an incision wound infection model that was recently established in the lab (45). In the peritonitis model, the Δ5Fe strain showed ∼1-log reduction (*p*≤0.0001) in the number of total bacteria recovered from the peritoneal cavity 48 hours post-infection when compared to the parent, Δ*fitAB*, Δ*emtB* and Δ*fitAB*Δ*emtB* strains (Fig. 8C). However, parent and all mutants, including Δ5Fe, were recovered in similar numbers from spleens (Fig. 8C). On the other hand, with exception of Δ*emtB*, all mutants were recovered from wounds in significantly lower numbers (*p*≤0.05) when compared to wounds infected with the parent strain (Fig. 8D).

### *E. faecalis* can utilize heme as an iron source for *E. faecalis*

To this point, the results obtained indicate that *E. faecalis* relies on the cooperative activity of at least five iron uptake systems to overcome iron deficiency. However, the *in vivo* results suggest that *E. faecalis* can deploy additional strategies to quench its need for iron during infection. Because the most abundant source of iron in mammals is in the form of heme whereby an iron ion is coordinated to a porphyrin molecule, and considering that some of the most successful invasive pathogens encode at least one dedicated heme import systems (12, 26-29, 46-49), we suspected that *E. faecalis* can also use heme as an iron source. In fact, *E. faecalis* has at least two heme-dependent enzymes, catalase (KatA) and cytochrome oxidase (CydAB) (50-52), and a heme exporter (HrtAB) and heme-sensing regulator (FhtR) that are used to overcome heme intoxication (53). Yet, *E. faecalis* does not encode the machinery for heme biosynthesis or systems homologous to any of the more conserved heme uptake systems, such that it remains elusive how *E. faecalis* acquires extracellular heme. Next, we asked if supplementation of the growth media with 10 μM heme could restore growth of the Δ5Fe strain in LM-FMC[-Fe]. As suspected, the addition of heme greatly increased growth rates and yields of the Δ5Fe strain in iron-depleted media albeit it also enhanced the final growth yield of the parent strain (Fig. 9A). Most likely, the beneficial effects of heme on cell growth are due to heme serving as the enzymatic co-factor for cytochrome oxidase and an iron source. To further probe the role of heme in iron homeostasis, we compared intracellular levels of heme and iron in parent and Δ5Fe strains grown in LM-FMC[-Fe] plus or minus 10 μM heme. As expected, heme was undetectable unless it was added to the growth media with both strains accumulating comparable levels of heme when grown in heme-supplemented LM-FMC (Fig. 9B). Importantly, heme supplementation more than doubled intracellular iron levels in the parent strain and restored intracellular iron homeostasis in the Δ5Fe strain (Fig. 9C). Collectively, these results reveal that E. faecalis can internalize heme and then degrade intracellularly to release the iron ion. On a separate note, the differences in intracellular iron levels in the Δ5Fe strain grown in LM-FMC[-Fe] that were significantly lower but quantifiable in Fig. 7A but below the limit of detection in Fig. 9C are a faithful representation of the variations that we observe between differences batches of media.

**FIG 9.**
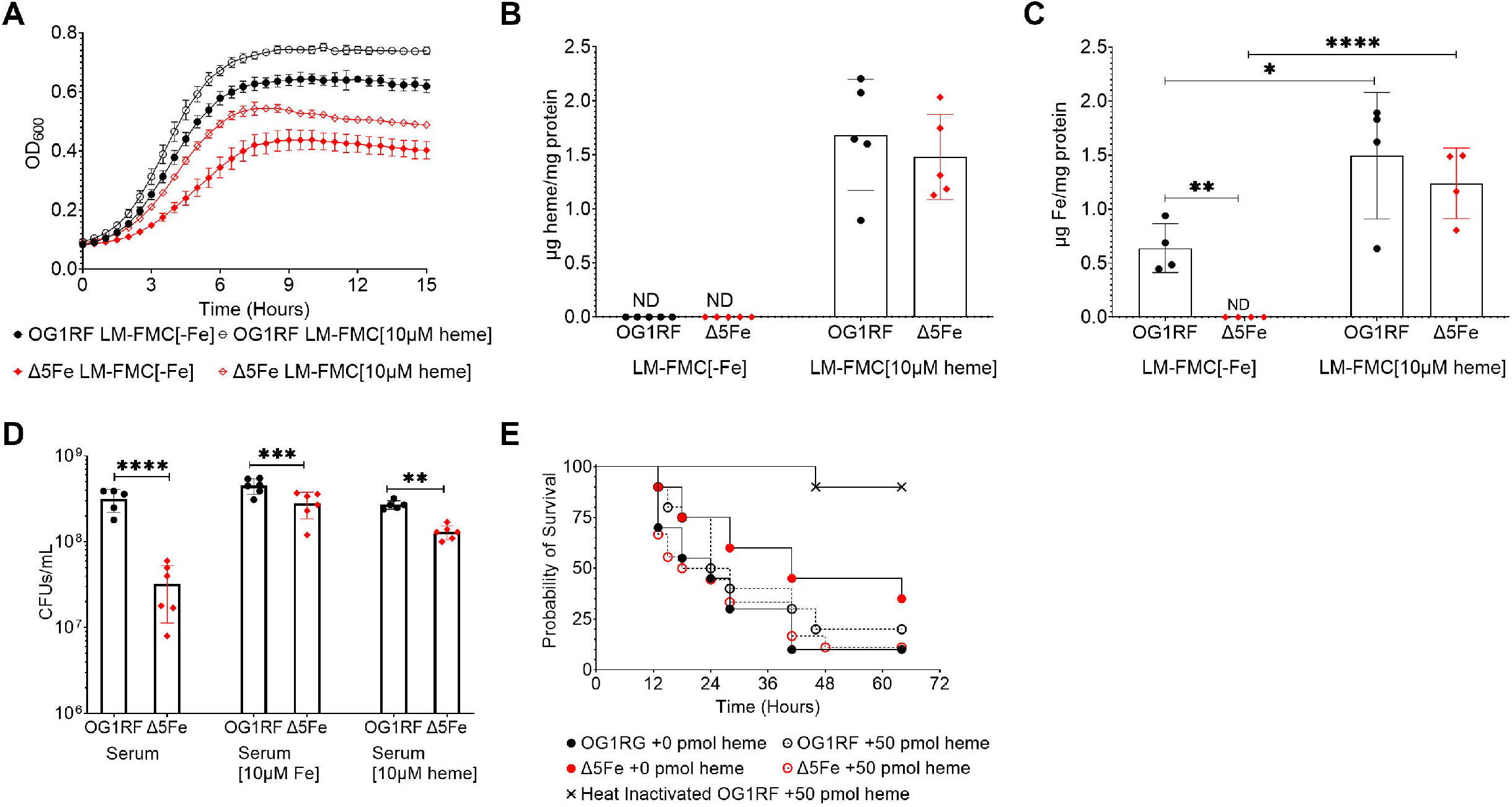
Heme restores growth and virulence of the *E. faecalis* Δ5Fe strain in iron depleted environments. (A) Growth of strains OG1RF or Δ5Fe in LM-FMC[-Fe] with or without 10 μM heme supplementation. (B) Intracellular heme content determined from cultures grown to mid-log phase in LM-FMC[-Fe] with or without 10 μM heme supplementation. Absorbances of and chloroform extract samples and heme standards were used in the correction equation Ac = 2 × *A*_388_ − (*A*_450_ + *A*_330_) and normalized to total protein content. (C) ICP-OES analysis of intracellular iron content of OG1RF and Δ5Fe strains grown in LM-FMC[-Fe] with or without heme supplementation. (B-C) Significance was determined by two-way ANOVA and a Dunnett’s post comparison test. (D) 24-hours growth of OG1RF and Δ5Fe in fresh human serum with or without supplementation with 10 μM iron or 10 μM heme. The experiment was performed on two separate occasions with three bacterial biological replicates. Error bars denote SEM and significance was obtained using a 2way ANOVA with a Sidak’s multiple comparison test. (E) Larvae of *G. mellonella* were injected with 50 pmol of heme or PBS 1 hour prior to infection with the OG1RF or Δ5Fe strains. Control group was injected with heat-killed (HK) OG1RF. Kaplan-Meyer curve shown is representative of six independent experiments with 20 larvae per experiment. Significance was determine using the Mantel-Cox log-rank test. *p≤0.05, **p≤0.01, ***p≤0.001, and ****p≤0.0001.

Next, we asked if exogenous heme could restore growth of Δ5Fe in human sera or its virulence in the *G. mellonella* model. While the fresh human sera used is expected to contain iron-sequestering and heme-sequestering proteins, addition of 10 μM FeSO_4_ or 10 μM heme to the sera rescued the growth defect phenotype of the Δ5Fe strain without providing a noticeable growth advantage to the parent strain (Fig 9D). Because oxygen is not transported via hemoglobin/Fe-heme complexes in insects but rather through binding to two copper ions coordinated by histidine residues in hemocyanins (54), non-hematophagous insects such as *G. mellonella* are considered to be heme-free (55). Thus, in the last set of experiments, we injected the hemolymph of *G. mellonella* with 50 pmol heme *b* (in the form of hemin) one hour prior to infecting the larvae with the desired *E. faecalis* strain. While heme administration did not affect the pathogenic behavior of the parent strain, it fully restored virulence of Δ5Fe strain (Fig. 9E). These results led us to conclude that *E. faecalis* can acquire heme from the environment and that host-derived heme is an important source of iron during infection.

## DISCUSSION

Despite the nearly universal role of iron in host-pathogen interactions, (6, 7, 14, 19-23, 56), very little is currently known about the mechanisms utilized by *E. faecalis* to obtain iron from the extracellular milieu and much less so about the contribution of iron import systems to enterococcal fitness and pathogenic behavior. In a series of studies that spanned through two decades, Lisiecki and colleagues were the first to propose that enterococci utilize multiple strategies to scavenge iron, which included production of siderophores, expression of high-affinity iron transporters, and an undefined capacity to seize iron directly from host transferrin and lactoferrin (57-59). Yet, most of their observations have not been validated by others and, at least in the case of siderophore production, appears to be incorrect based on the absence of the machinery necessary for siderophore biosynthesis in *E. faecalis* genomes. Indeed, our multiple attempts to detect siderophore production in different strains of *E. faecalis* or *E. faecium* using the CAS (Chrom Azurol S) method (60) were not successful (data not shown). In addition to the work by Lisiecki and colleagues, *in silico* and transcriptome-based analyses using a Δ*fur* mutant have indicated that *E. faecalis* possess three highly conserved iron import systems, the ferrous iron transporter FeoAB, the ferrichrome transporter FhuDCBG, and the dual iron/manganese transporter EfaCBA (31, 33-35).

In this report, we validated previous studies (61) showing that either *E. faecalis* or *E. faecium* isolates can grow in media that can be considered virtually iron-free (0 to 0.003 parts per million iron depending on the batch of media). While the remarkable capacity of enterococci and of other lactic acid bacteria to grow under nearly iron-free conditions has been attributed to their “manganese-centric” nature, intracellular iron quantifications revealed that *E. faecalis* accumulates similar amounts of iron when grown in iron replete or iron depleted media. Rather than suggesting that *E. faecalis* does not require iron for growth as once suggested (62), we believe that iron is such an essential micronutrient to *E. faecalis* that it evolved multiple, diverse, and highly efficient systems to acquire and maintain iron homeostasis.

In addition to the conserved iron import systems EfaCBA, FeoAB and FhuDCBG, our transcriptomic analysis identified two novel ABC-type iron transporters that were named FitABCD and EmtABC. While this is the first time that EmtABCD is linked to iron uptake, FitABCD was previously shown to be a member of the Fur regulon (33). Moreover, *ex vivo* and *in vivo* transcriptome analysis have shown that, except for *fhuDCBG*, all other systems are highly expressed under physiologically relevant conditions. For example, *fitABCD* was upregulated by ∼4-fold in both human blood and human urine *ex vivo*, and 23- to 42-fold in a subdermal abscess rabbit model (63-65). The dual iron/manganese transporter *efaCBA* was upregulated ∼3-fold in either human blood or urine (63, 64), ∼2-fold in the abscess rabbit model (65), and ∼7-fold in a peritonitis mouse model (44). Finally, *emtABC* was upregulated ∼3-fold in human blood (63) and *feoAB* upregulated by ∼2-fold in human urine and ∼5-fold in the abscess rabbit model (64, 65). In this study, we showed that the individual responses of these transcriptional units to iron depletion can be divided into early (*efaCBA* and *emtABC*) and late (*fitABCD, feoAB*, and *fhuDCBG*) responders. Moreover, all late responders have been shown to be regulated by Fur (33) while *efaCBA* is regulated by EfaR (38, 66). Through bioinformatic analysis, we identified a putative EfaR-binding motif (38) located 13-bp upstream from the *emtABC* translational start site. Therefore, it is conceivable that transcriptional induction of iron acquisition systems is distinctly controlled by Fur and EfaR. The occurrence of these two distinct transcriptional surges is reminiscent of the stepwise induction of iron uptake systems in *B. subtilis* whereby elemental iron, ferric citrate, and petrobactin operons are induced in the first wave and bacillibactin synthesis and uptake, and hydroxamate siderophore uptake induced in the second wave (67). However, in *B. subtilis*, this sequential activation was solely dependent on the Fur regulator with subsequent experiments demonstrating that the stepwise transcriptional activation correlated with Fur operator occupancy *in vivo* (67). More studies are needed to determine if EfaR directly regulates *emtABC* and to validate the working hypothesis that iron starvation responses in *E. faecalis* can be separated by EfaR-regulated early responders and Fur-regulated late responders.

Even though systems homologous to EfaABC, FeoAB and FhuDCBG are widespread and have been relatively well characterized in bacteria (21, 56, 68-71), predicted proteins sharing high levels (≥80%) of similarity with FitABCD or EmtABC are almost entirely restricted to species of the enterococcacea family, with FitD and EmtC sharing slightly lower similarity (∼60-65%) with substrate-binding proteins from selected streptococci and bacilli. Of interest, the *B. subtilis* YclNOPQ transporter is responsible for uptake of the petrobactin siderophore (37), raising the possibility that FitABC mediates siderophore uptake. This might also be the case of FhuDCBG that mediates uptake of ferric hydroxamate-type siderophores in other bacteria (72). As mentioned above, it appears that enterococci cannot synthesize its own siderophores such that these systems might be involved in xenosiderophore uptake or other types of iron source. Additional studies are necessary to determine the iron species specificity and affinities of FitABCD and EmtABC.

The finding that *E. faecalis* possesses multiple systems to acquire iron is not surprising when considering their capacity to inhabit a variety of niches within the host, from the gastrointestinal tract to the skin, oral cavity, and the genitourinary tract, and to remain viable for prolonged periods when excreted into the environment. In addition, there are numerous examples in the literature describing how bacteria deploy multiple and complementary strategies to maintain iron homeostasis. As mentioned above, *B. subtilis* encodes transporters for the uptake of elemental iron, ferric citrate, different types of siderophores, in addition to producing its own siderophore and cognate import system (67). Similarly, *S. aureus* encodes transporters for elemental iron, iron hydroxamates, and synthesizes two types of siderophores (staphyloferrin A and B) along with their cognate importers (21). In addition, *S. aureus* encodes the Isd system that mediates binding, degradation, and uptake of iron-heme complexes (49, 73). Similar to *S. aureus*, some of the major pathogenic species of streptococci encode a suite of elemental iron, siderophore and heme transport systems (21, 29, 74, 75). As expected, inactivation of a single iron transport system had minimal or no impact on the ability of *E. faecalis* to grow under severe iron deficiency. To demonstrate this functional overlap, we generated a quintuple (Δ5Fe) mutant lacking all five systems. The Δ5Fe strain grew poorly in media without an added iron source, accumulated considerably less intracellular iron than the parental strain, and showed major deficiency in elemental iron uptake. The Δ5Fe strain also failed to grow in media depleted of both iron and manganese, likely because EfaCBA is a dual iron and manganese transporter. Despite these observations and considering that vertebrate hosts actively restrict both iron and manganese during infection, we found that the virulence potential of Δ5Fe varied depending on the model used and, possibly, the site of infection within the vertebrate host. While virulence of Δ5Fe was markedly attenuated in *G. mellonella*, and the mutant was recovered in significantly lower numbers from mouse peritoneal cavity and infected mouse wounds, parent and Δ5Fe strains were recovered in similar numbers from spleens in the peritonitis model. We suspected that the capacity to utilize heme as an iron source was behind this apparent conflicting result. To explore this possibility, we conducted a series of experiments that showed that *E. faecalis* is indeed capable of using heme as an iron source and that heme supplementation restores virulence of the Δ5Fe strain in *G. mellonella*. While *E. faecalis* does not possess the machinery for heme biosynthesis and does not require heme for growth (52, 76), it encodes at least two heme-dependent enzymes, cytochrome *bd* oxidase and catalase, such that it must have the capacity to obtain heme from the extracellular milieu. Yet, systems homologous to known heme transport systems such as the *S. aureus* Isd or the *S. pyogenes* Sia are absent in enterococcal genomes. During preparation of this manuscript, the Kline lab provided initial evidence that the ABC-type integral membrane proteins CydCD, previously implicated in cytochrome assembly and cysteine export (77, 78), mediate heme uptake (79). While additional studies are needed to confirm the role of CydCD in heme uptake, it is also apparent that CydCD are not working alone in heme uptake since heme-dependent catalase activity can be still detected in *cydABCD* mutants (51). Studies to identify the elusive heme import systems of *E. faecalis*, to separate the significance of heme as a nutrient and as an iron source, and to determine how disruption of heme uptake will affect the pathogenic potential of *E. faecalis* in different types of infection are ongoing.

## MATERIALS AND METHODS

### Bacterial strains and growth conditions

Bacterial strains used in this study are listed in Table 2. All *E. faecalis* strains were routinely grown aerobically at 37**°**C in brain heart infusion (Difco). For controlled growth under metal-depleted conditions, we used the chemically defined FMC media originally developed for cultivation of oral streptococci (36), with minor modifications. Specifically, the base media was prepared without any of the metal components (magnesium, calcium, iron, and manganese) and treated with Chelex (BioRad) to remove contaminating metals. The pH was adjusted to 7.0 and filter sterilized. Magnesium and calcium solutions were prepared using National Exposure Research Laboratory (NERL) trace metal grade water, filter sterilized, and then added to the media. Iron and manganese solutions were also prepared using NERL trace metal grade water, filter sterilized, and added to the media as indicated in the text and figure legends. For RNA-seq analysis, overnight BHI cultures of *E. faecalis* OG1RF were diluted 1:100 in FMC[+Fe] or FMC[-Fe] and grown to an OD_600_ of 0.5 before cells were collected for RNA isolation. For reverse transcriptase quantitative PCR (RT-qPCR) analysis, RNA was isolated from cells grown in FMC[+Fe] and then shifted to FMC[-Fe] with aliquots taken 10, 30, and 60 minutes after the shift. To generate growth curves, cultures were grown in BHI to an OD_600_ of 0.25 (early exponential phase) and then diluted 1:200 into fresh media that were either BHI, FMC or LM-FMC supplemented with heme, iron, and/or manganese as indicated in the text and figure legends. Cell growth was monitored using the Bioscreen growth reader (Oy Growth Curves).

**TABLE 2.**
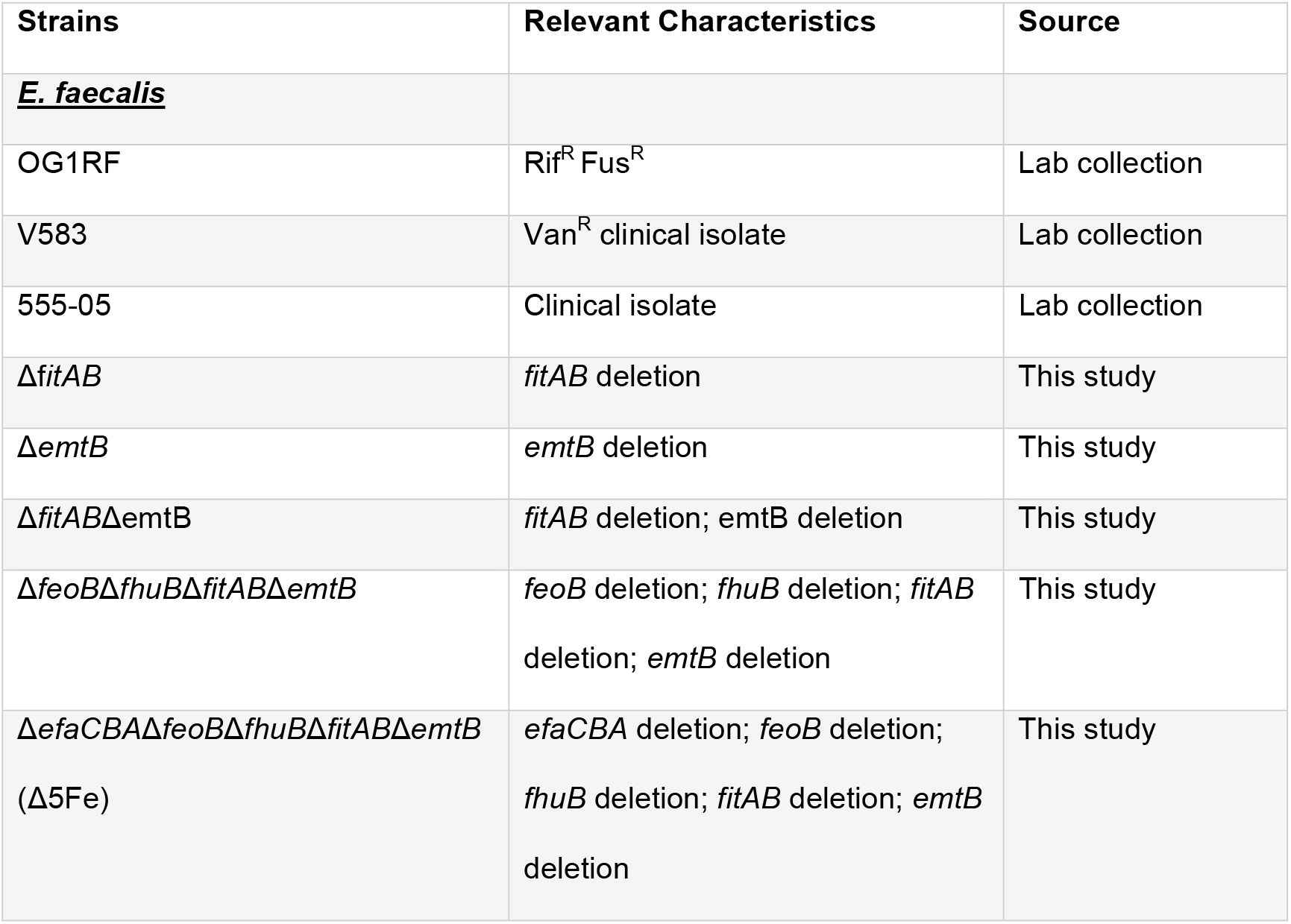
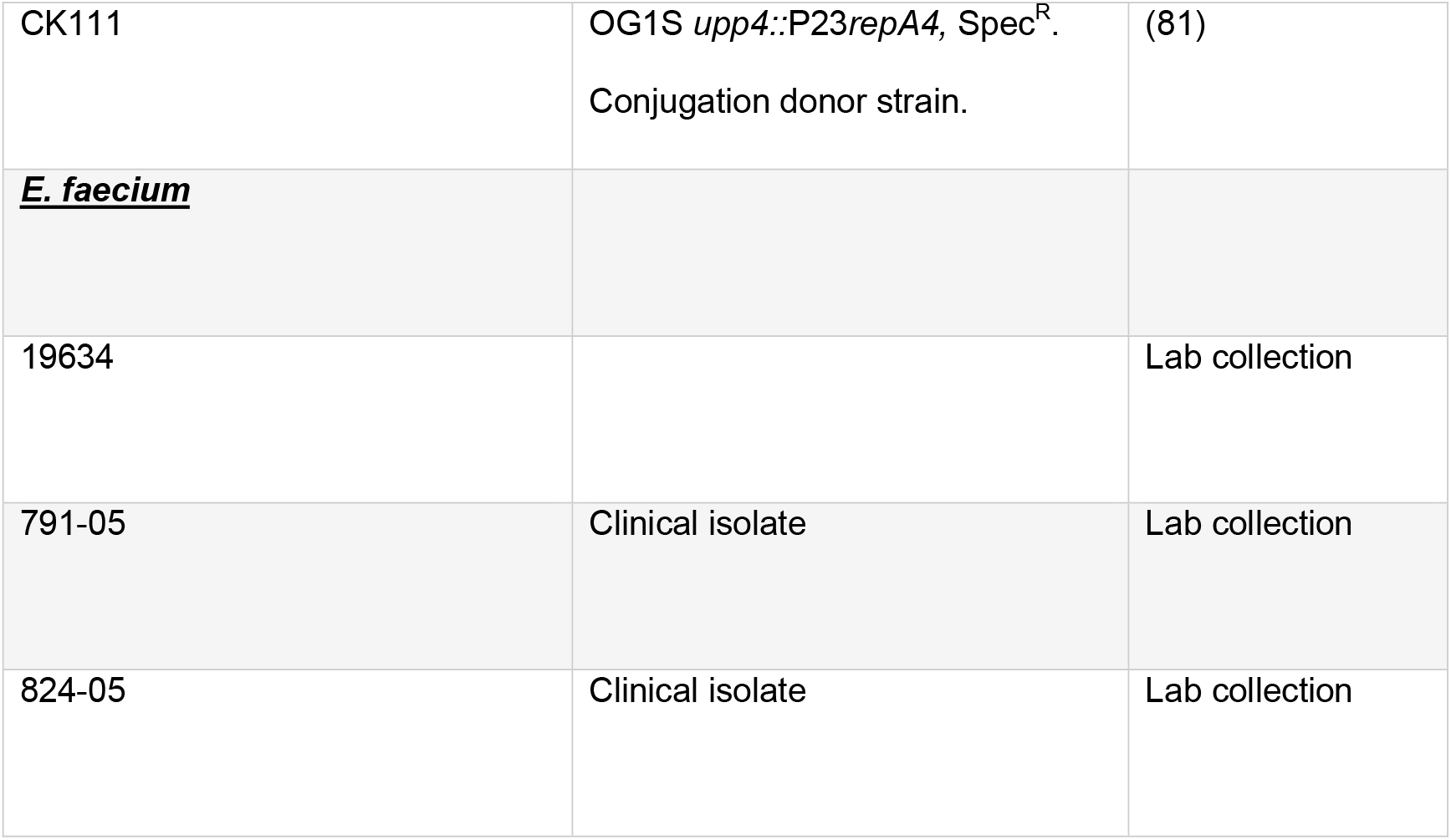
Strains of *E. faecalis* and *E. faecium* used in this study.

### Construction of mutant strains

Markerless deletions of *fitAB, emtB, feoB* or *fhuB* in *E. faecalis* OG1RF strain was carried out using the pCJK47 genetic exchange system (31). Briefly, PCR products with ∼1 kb in size flanking each coding sequence were amplified with the primers listed in Table S3. To avoid unanticipated polar effects, amplicons included either the first or last residues of the coding sequences. Cloning of amplicons into the pCJK47 vector, electroporation, and conjugation into *E. faecalis* strains and isolation of single mutant strains (Δ*fitAB*, Δ*emtB*, Δ*feoB* and Δ*fhuB*) were carried out as previously described (31). The Δ*fitAB*Δ*emtB* double mutant was obtained by conjugating the pCJK-*emtB* plasmid into the Δ*fitAB* mutant. Then, a triple mutant was obtained by conjugating the pCJK-*fhuB* plasmid into the Δ*fitAB*Δ*emtB* double mutant and a quadruple obtained by conjugation of pCJK47-*feoB* into the Δ*fitAB*Δ*emtB*Δ*fhuB* triple mutant. Finally, the quintuple mutant was isolated by conjugation of pCJK-*efaCBA* (31) into the quadruple mutant. All gene deletions were confirmed by PCR sequencing of the insertion site and flanking region.

### RNA analysis

Total RNA was isolated from *E. faecalis* OG1RF cells grown to mid-log phase in FMC[+Fe] or FMC[-Fe] or grown to mid-log phase in FMC[+Fe] and transferred to FMC[-Fe] following the methods described elsewhere (80). The RNA was precipitated with ice-cold isopropanol and 3 M sodium acetate (pH 5) at 4°C before RNA pellets were suspended in nuclease-free H_2_O and treated with DNase I (Ambion) for 30 min at 37°C. Then, ∼ 100 μg of RNA per sample was further purified using the RNeasy kit (Qiagen), which includes a second on-column DNase digestion. Sample quality and quantity were assessed on an Agilent 2100 Bioanalyzer at the University of Florida Interdisciplinary Center for Biotechnology Research (UF-ICBR). Messenger RNA (5 μg total RNA per sample) was enriched using a MICROBExpress bacterial mRNA purification kit (Thermo Fisher) and cDNA libraries containing unique barcodes generated from 100 ng mRNA using the Next UltraII Directional RNA Library Prep kit for Illumina (New England Biolabs). The individual cDNA libraries were assessed for quality and quantity by Qubit, diluted to 10 nM each and equimolar amounts of cDNA pooled together. The pooled cDNA libraries were subjected to deep sequencing at the UF-ICBR using the Illumina NextSeq 500 platform. Read mapping was performed on a Galaxy server hosted by the University of Florida Research Computer using Map with Bowtie for Illumina and the *E. faecalis* OG1RF genome (GenBank accession no. NC_017316.1) used as reference. The reads per open reading frame were tabulated with htseq-count. Final comparisons between bacteria grown in FMC[+Fe] and FMC[-Fe] were performed with Degust (http://degust.erc.monash.edu/), with a false-discovery rate (FDR) of 0.05 and after applying a 2-fold change cutoff.

### ICP-OES

Trace metal content in bacteria or growth media was determined by inductively coupled plasma-optical emission spectrometry (ICP-OES). For quantification of trace metals in the different media used, 18 ml of prepared media (BHI, FMC or LM-FMC) were digested with 2 ml trace-metal grade 35% HNO_3_ at 90°C for 1 hour. For intracellular metal quantification, cell pellets from overnight BHI cultures were washed once in 0.5 mM EDTA and twice in trace-metal grade PBS to remove extracellular metals and diluted 1:50 in LM-FMC with or without iron or heme supplementation as described in the results section. Cultures were grown aerobically at 37°C to an OD_600_ 0.4, the cell pellets collected by centrifugation, washed once in 0.5 mM EDTA and twice in trace metal grade PBS to remove extracellular metals. A 10 ml aliquot of resuspended cell pellet was saved for total protein quantification and 40 ml of the suspension used for metal quantification. For this, cell suspensions were digested in 2 ml 35% HNO_3_ at 90°C for 1 hour, and the digested suspension diluted 1:10 in reagent-grade H_2_O. Metal content was determined using a 5300DV ICP Atomic Emission Spectrometer (Perkin Elmer) at the University of Florida Institute of Food and Agricultural Sciences Analytical Services Laboratories, and the data normalized to total protein content that was determined by the bicinchoninic acid (BCA) assay (Sigma).

### ^55^Fe uptake

For ^55^Fe uptake experiments, nitrocellulose membranes were pre-wet in 1 M NiSO_4_ solution to prevent nonspecific binding of ^55^Fe (Perkin-Elmer) to the membranes. Overnight cultures of *E. faecalis* parent and mutant strains grown in LM-FMC[-Fe] were diluted 30-fold in LM-FMC[-Fe], and grown to mid-log phase (OD_600_ ∼0.5), at which point 10 μM ^55^Fe was added to each culture and incubated at 37°C. At 0, 15, 30, and 60 minutes, 200 μl aliquots were transferred to the pre-wet nitrocellulose membrane placed in a slot blot apparatus. Free ^55^Fe was removed by four washes with 100 mM sodium citrate buffer using vacuum filtration. The membranes were air dried, cut, and dissolved in 4 ml scintillation counter cocktail. Radioactivity was measured by scintillation with “wide open” window setting using a Beckmann LSC6000 scintillation counter. The count per million (cpm) values from ^55^Fe free cells were obtained and subtracted from the cpm of treated cells. The efficiency of the machine was ∼30.8% and was used to convert cpm to disintegrations per minute (dpm), which was then converted to molarity and normalized to CFU.

### Intracellular heme quantification

Cultures were grown under the same conditions used for trace metal quantifications by ICP-OES. After washing in trace-metal grade PBS, pellets were suspended in 1ml DMSO and lysed using a bead beater. Cellular heme was determined using the acidified chloroform extraction method following the protocols detailed elsewhere (29). Absorbance of the organic phases at 388, 450, and 330 nm were determined using a GENESYS™ 30 Visible Spectrophotometer (ThermoScientific™). Heme content was determined by plugging absor− bance values of samples and heme standards into the correction equation Ac = 2 × *A*_388_ − (*A*_450_ + *A*_330_) and were normalized by total protein content.

### Growth in human serum

Blood from B^+^ healthy donors was obtained from LifeSouth Community Blood Centers in Gainesville, Florida (IRB 202100899). Each experiment was performed with pooled serum isolated from blood of 3 individual donors. Where indicated, serum was supplemented with 10μM FeSO_4_ or 10 μM heme. After overnight incubation in BHI at 37°C, cell pellets were collected, washed once in 0.5 mM EDTA in trace metal grade PBS, twice in trace metal grade PBS, and sub-cultured into serum at ∼1.5 × 10^6^ CFU ml^−1^ with constant rotation at 37°C. Total CFU at selected intervals was determined by serial dilution and plated on tryptic soy agar (TSA) containing 200 μg ml ^−1^ rifampicin and 10 μg ml^−1^ fusidic acid.

### *Galleria mellonella* infection

Larvae of *G. mellonella* was used to assess virulence of parent and selected mutants as previously described (31). Briefly, groups of 20 larvae (200–300 mg in weight) were injected with 5 μl of bacterial inoculum containing ∼5 × 10^5^ CFU. To investigate the impact of exogenous heme supplementation, larvae were injected with either trace metal grade PBS or 50 pmol heme one hour prior to infection. Larvae injected with heat-inactivated *E. faecalis* OG1RF (30 min at 100°C), 50 pmol heme, or PBS were used as controls. After infection, larvae were kept at 37°C and their survival monitored for up to 96 hours.

### Mouse intraperitoneal infection

These experiments were performed under protocol 202200000241 approved by the University of Florida Institutional Animal Care and Use Committee (IACUC). The mouse peritonitis infection model has been described previously (43) such that only a brief overview of the model is provided below. To prepare the bacterial inoculum, bacteria were grown in BHI to an OD_600_ of 0.5, the cells pellets collected, washed once in 0.5 mM EDTA and twice in trace metal grade PBS, and suspended in PBS at ∼2 × 10^8^ CFU ml^−1^. Seven-week-old C57BL6J mice purchased from Jackson laboratories were intraperitoneally injected with 1 ml of bacterial suspension and euthanized by CO_2_ asphyxiation 48-h post-infection. The abdomen was opened to expose the peritoneal lining, 5 ml of cold PBS injected into the peritoneal cavity with 4 ml retrieved as the peritoneal wash. Quantification of bacteria within the peritoneal wash was determined by plating serial dilutions on TSA containing 200 μg ml ^−1^ rifampicin and 10 μg ml^−1^ fusidic acid. For bacterial enumeration inside spleens, spleens were surgically removed, briefly washed in 70% ethanol followed by rinsing in sterile PBS, homogenized in 1 ml PBS, serially diluted, and plated on selective TSA plates.

### Mouse wound infection

These experiments were performed under protocol 202011154 approved by the University of Florida IACUC. The bacterial inoculum was prepared as described for the peritonitis model, but cell pellets were concentrated to 1 × 10^10^ CFU ml^−1^ and stored on ice until infection. Seven-week old C57BL6J mice purchased from Jackson laboratories were anesthetized using isoflurane, their backs shaved, and the incision wound created using a 6mm biopsy punch. Wounds were infected with 10 μl of culture and covered with Tegaderm™ dressing. 72 hours post infection, mice were euthanized by CO_2_ asphyxiation, the wounds were excised, and the wounds homogenized in 1 ml PBS. The wound homogenates were serially diluted and plated on selective TSA plates.

## Acknowledgements

This study was supported by NIH-NIAID R21 AI137446 to J.A.L. D.N.B. was supported by NIH-NIDCR Training Grant T90 DE021990 and by American Heart Association Pre-doctoral Fellowship 907592.

## Data availability

Gene expression data have been deposited in the NCBI Gene Expression Omnibus (GEO) database (https://www.ncbi.nlm.nih.gov/geo). The GEO Series accession number is pending.

## SUPPLEMENTAL FIGURE LEGENDS

**FIG S1** Growth of OG1RF, Δ*feoB*, Δ*fhuB*, and Δ*efaCBA* in (A) BHI, (B) LM-FMC[+Fe], (C) LM-FMC[-Fe], and (D) LM-FMC[-Fe/-Mn]. Growth was monitored by measuring OD_600_ every 30 minutes using an automated growth reader. Error bars denote standard deviations from three biological replicates.

**FIG S2** Growth of Δ*fitAB*Δ*emtB*, and Δ*feoB*Δ*fhuB*ΔfitABΔemtB in (A) LM-FMC[+Fe] and (B) LM-FMC[-Fe]. Growth was monitored by measuring OD_600_ every 30 minutes using an automated growth reader. Error bars denote standard deviations from three biological replicates.

**FIG S3** 24-hours growth of OG1RF, Δ*fitAB*, Δ*emtB*, Δ*fitAB*Δ*emtB*, and Δ5Fe in fresh human serum with. The experiment was performed on two separate occasions with three bacterial biological replicates. Error bars denote SEM and significance was obtained using a one-way ANOVA with a Holm-Šídák’s multiple comparisons test.

## REFERENCES

1. Magill SS, O’Leary E, Edwards JR, Emerging Infections Program Healthcare-Associated I, Antimicrobial Use Hospital Prevalence Survey T. Changes in Prevalence of Health Care-Associated Infections. Reply. N Engl J Med. 2019;380(11):1085–6. Epub 2019/03/14. doi: 10.1056/NEJMc1817140. PubMed PMID: 30865811; PMCID: PMC7976444.

2. Solomon SL, Oliver KB. Antibiotic resistance threats in the United States: stepping back from the brink. Am Fam Physician. 2014;89(12):938–41. Epub 2014/08/28. PubMed PMID: 25162160.

3. Arias CA, Murray BE. The rise of the Enterococcus: beyond vancomycin resistance. Nat Rev Microbiol. 2012;10(4):266–78. Epub 2012/03/17. doi: 10.1038/nrmicro2761. PubMed PMID: 22421879; PMCID: PMC3621121.

4. Gaca AO, Lemos JA. Adaptation to Adversity: the Intermingling of Stress Tolerance and Pathogenesis in Enterococci. Microbiol Mol Biol Rev. 2019;83(3). Epub 2019/07/19. doi: 10.1128/MMBR.00008-19. PubMed PMID: 31315902; PMCID: PMC6710459.

5. Kao PHN, Kline KA. Dr. Jekyll and Mr. Hide: How Enterococcus faecalis Subverts the Host Immune Response to Cause Infection. J Mol Biol. 2019;431(16):2932–45. Epub 2019/05/28. doi: 10.1016/j.jmb.2019.05.030. PubMed PMID: 31132360.

6. Begg SL. The role of metal ions in the virulence and viability of bacterial pathogens. Biochem Soc Trans. 2019;47(1):77–87. Epub 2019/01/11. doi: 10.1042/BST20180275. PubMed PMID: 30626704.

7. Chandrangsu P, Rensing C, Helmann JD. Metal homeostasis and resistance in bacteria. Nat Rev Microbiol. 2017;15(6):338–50. Epub 2017/03/28. doi: 10.1038/nrmicro.2017.15. PubMed PMID: 28344348; PMCID: PMC5963929.

8. Becker KW, Skaar EP. Metal limitation and toxicity at the interface between host and pathogen. FEMS Microbiol Rev. 2014;38(6):1235–49. Epub 2014/09/12. doi: 10.1111/1574-6976.12087. PubMed PMID: 25211180; PMCID: PMC4227937.

9. Kehl-Fie TE, Skaar EP. Nutritional immunity beyond iron: a role for manganese and zinc. Curr Opin Chem Biol. 2010;14(2):218–24. Epub 2009/12/18. doi: 10.1016/j.cbpa.2009.11.008. PubMed PMID: 20015678; PMCID: PMC2847644.

10. Nairz M, Dichtl S, Schroll A, Haschka D, Tymoszuk P, Theurl I, Weiss G. Iron and innate antimicrobial immunity-Depriving the pathogen, defending the host. J Trace Elem Med Biol. 2018;48:118–33. Epub 2018/05/19. doi: 10.1016/j.jtemb.2018.03.007. PubMed PMID: 29773170.

11. Dev S, Babitt JL. Overview of iron metabolism in health and disease. Hemodial Int. 2017;21 Suppl 1:S6–S20. Epub 2017/03/16. doi: 10.1111/hdi.12542. PubMed PMID: 28296010; PMCID: PMC5977983.

12. Pantopoulos K, Porwal SK, Tartakoff A, Devireddy L. Mechanisms of mammalian iron homeostasis. Biochemistry. 2012;51(29):5705–24. Epub 2012/06/19. doi: 10.1021/bi300752r. PubMed PMID: 22703180; PMCID: PMC3572738.

13. Vogt AS, Arsiwala T, Mohsen M, Vogel M, Manolova V, Bachmann MF. On Iron Metabolism and Its Regulation. Int J Mol Sci. 2021;22(9). Epub 2021/05/01. doi: 10.3390/ijms22094591. PubMed PMID: 33925597; PMCID: PMC8123811.

14. Cassat JE, Skaar EP. Iron in infection and immunity. Cell Host Microbe. 2013;13(5):509–19. Epub 2013/05/21. doi: 10.1016/j.chom.2013.04.010. PubMed PMID: 23684303; PMCID: PMC3676888.

15. Wang J, Pantopoulos K. Regulation of cellular iron metabolism. Biochem J. 2011;434(3):365–81. Epub 2011/02/26. doi: 10.1042/BJ20101825. PubMed PMID: 21348856; PMCID: PMC3048577.

16. Hao L, Shan Q, Wei J, Ma F, Sun P. Lactoferrin: Major Physiological Functions and Applications. Curr Protein Pept Sci. 2019;20(2):139–44. Epub 2018/05/15. doi: 10.2174/1389203719666180514150921. PubMed PMID: 29756573.

17. Zygiel EM, Nolan EM. Exploring Iron Withholding by the Innate Immune Protein Human Calprotectin. Acc Chem Res. 2019;52(8):2301–8. Epub 2019/08/06. doi: 10.1021/acs.accounts.9b00250. PubMed PMID: 31381301; PMCID: PMC6702060.

18. Lopez CA, Skaar EP. The Impact of Dietary Transition Metals on Host-Bacterial Interactions. Cell Host Microbe. 2018;23(6):737–48. Epub 2018/06/15. doi: 10.1016/j.chom.2018.05.008. PubMed PMID: 29902439; PMCID: PMC6007885.

19. Ratledge C, Dover LG. Iron metabolism in pathogenic bacteria. Annu Rev Microbiol. 2000;54:881–941. doi: DOI 10.1146/annurev.micro.54.1.881. PubMed PMID: WOS:000165272300028.

20. Braun V, Hantke K. Recent insights into iron import by bacteria. Curr Opin Chem Biol. 2011;15(2):328–34. Epub 2011/02/01. doi: 10.1016/j.cbpa.2011.01.005. PubMed PMID: 21277822.

21. Sheldon JR, Heinrichs DE. Recent developments in understanding the iron acquisition strategies of gram positive pathogens. FEMS Microbiol Rev. 2015;39(4):592–630. Epub 2015/04/12. doi: 10.1093/femsre/fuv009. PubMed PMID: 25862688.

22. Palmer LD, Skaar EP. Transition Metals and Virulence in Bacteria. Annu Rev Genet. 2016;50:67–91. Epub 2016/09/13. doi: 10.1146/annurev-genet-120215-035146. PubMed PMID: 27617971; PMCID: PMC5125913.

23. Nairz M, Haschka D, Demetz E, Weiss G. Iron at the interface of immunity and infection. Front Pharmacol. 2014;5:152. Epub 2014/08/01. doi: 10.3389/fphar.2014.00152. PubMed PMID: 25076907; PMCID: PMC4100575.

24. Wilson BR, Bogdan AR, Miyazawa M, Hashimoto K, Tsuji Y. Siderophores in Iron Metabolism: From Mechanism to Therapy Potential. Trends Mol Med. 2016;22(12):1077–90. Epub 2016/11/09. doi: 10.1016/j.molmed.2016.10.005. PubMed PMID: 27825668; PMCID: PMC5135587.

25. Cook-Libin S, Sykes EME, Kornelsen V, Kumar A. Iron Acquisition Mechanisms and Their Role in the Virulence of Acinetobacter baumannii. Infect Immun. 2022:e0022322. Epub 2022/09/07. doi: 10.1128/iai.00223-22. PubMed PMID: 36066263.

26. Jones CM, Niederweis M. Mycobacterium tuberculosis can utilize heme as an iron source. J Bacteriol. 2011;193(7):1767–70. Epub 2011/02/08. doi: 10.1128/JB.01312-10. PubMed PMID: 21296960; PMCID: PMC3067660.

27. Jochim A, Adolf L, Belikova D, Schilling NA, Setyawati I, Chin D, Meyers S, Verhamme P, Heinrichs DE, Slotboom DJ, Heilbronner S. An ECF-type transporter scavenges heme to overcome iron-limitation in Staphylococcus lugdunensis. Elife. 2020;9. Epub 2020/06/10. doi: 10.7554/eLife.57322. PubMed PMID: 32515736; PMCID: PMC7299338.

28. Contreras H, Chim N, Credali A, Goulding CW. Heme uptake in bacterial pathogens. Curr Opin Chem Biol. 2014;19:34–41. Epub 2014/05/02. doi: 10.1016/j.cbpa.2013.12.014. PubMed PMID: 24780277; PMCID: PMC4007353.

29. Chatterjee N, Cook LCC, Lyles KV, Nguyen HAT, Devlin DJ, Thomas LS, Eichenbaum Z. A Novel Heme Transporter from the Energy Coupling Factor Family Is Vital for Group A Streptococcus Colonization and Infections. J Bacteriol. 2020;202(14). Epub 2020/05/13. doi: 10.1128/JB.00205-20. PubMed PMID: 32393520; PMCID: PMC7317044.

30. Han R, Niu M, Liu S, Mao J, Yu Y, Du Y. The effect of siderophore virulence genes entB and ybtS on the virulence of Carbapenem-resistant Klebsiella pneumoniae. Microb Pathog. 2022;171:105746. Epub 2022/09/06. doi: 10.1016/j.micpath.2022.105746. PubMed PMID: 36064103.

31. Colomer-Winter C, Flores-Mireles AL, Baker SP, Frank KL, Lynch AJL, Hultgren SJ, Kitten T, Lemos JA. Manganese acquisition is essential for virulence of Enterococcus faecalis. PLoS Pathog. 2018;14(9):e1007102. Epub 2018/09/21. doi: 10.1371/journal.ppat.1007102. PubMed PMID: 30235334; PMCID: PMC6147510.

32. Lam LN, Brunson DN, Molina JJ, Flores-Mireles AL, Lemos JA. The AdcACB/AdcAII system is essential for zinc homeostasis and an important contributor of Enterococcus faecalis virulence. Virulence. 2022;13(1):592–608. Epub 2022/03/29. doi: 10.1080/21505594.2022.2056965. PubMed PMID: 35341449; PMCID: PMC8966984.

33. Latorre M, Quenti D, Travisany D, Singh KV, Murray BE, Maass A, Cambiazo V. The Role of Fur in the Transcriptional and Iron Homeostatic Response of Enterococcus faecalis. Front Microbiol. 2018;9:1580. Epub 2018/08/02. doi: 10.3389/fmicb.2018.01580. PubMed PMID: 30065712; PMCID: PMC6056675.

34. Lopez G, Latorre M, Reyes-Jara A, Cambiazo V, Gonzalez M. Transcriptomic response of Enterococcus faecalis to iron excess. Biometals. 2012;25(4):737–47. Epub 2012/03/27. doi: 10.1007/s10534-012-9539-5. PubMed PMID: 22447126.

35. Latorre M, Galloway-Pena J, Roh JH, Budinich M, Reyes-Jara A, Murray BE, Maass A, Gonzalez M. Enterococcus faecalis reconfigures its transcriptional regulatory network activation at different copper levels. Metallomics. 2014;6(3):572–81. Epub 2014/01/03. doi: 10.1039/c3mt00288h. PubMed PMID: 24382465; PMCID: PMC4131723.

36. Terleckyj B, Willett NP, Shockman GD. Growth of several cariogenic strains of oral streptococci in a chemically defined medium. Infect Immun. 1975;11(4):649–55. Epub 1975/04/01. doi: 10.1128/iai.11.4.649-655.1975. PubMed PMID: 1091546; PMCID: PMC415117.

37. Zawadzka AM, Kim Y, Maltseva N, Nichiporuk R, Fan Y, Joachimiak A, Raymond KN. Characterization of a Bacillus subtilis transporter for petrobactin, an anthrax stealth siderophore. Proc Natl Acad Sci U S A. 2009;106(51):21854–9. Epub 2009/12/04. doi: 10.1073/pnas.0904793106. PubMed PMID: 19955416; PMCID: PMC2799803.

38. Low YL, Jakubovics NS, Flatman JC, Jenkinson HF, Smith AW. Manganese-dependent regulation of the endocarditis-associated virulence factor EfaA of Enterococcus faecalis. J Med Microbiol. 2003;52(Pt 2):113–9. Epub 2003/01/25. doi: 10.1099/jmm.0.05039-0. PubMed PMID: 12543916.

39. Bergin D, Murphy L, Keenan J, Clynes M, Kavanagh K. Pre-exposure to yeast protects larvae of Galleria mellonella from a subsequent lethal infection by Candida albicans and is mediated by the increased expression of antimicrobial peptides. Microbes Infect. 2006;8(8):2105–12. Epub 2006/06/20. doi: 10.1016/j.micinf.2006.03.005. PubMed PMID: 16782387.

40. Kelly J, Kavanagh K. Caspofungin primes the immune response of the larvae of Galleria mellonella and induces a non-specific antimicrobial response. J Med Microbiol. 2011;60(Pt 2):189–96. Epub 2010/10/16. doi: 10.1099/jmm.0.025494-0. PubMed PMID: 20947665.

41. Zimmer DB, Eubanks JO, Ramakrishnan D, Criscitiello MF. Evolution of the S100 family of calcium sensor proteins. Cell Calcium. 2013;53(3):170–9. Epub 2012/12/19. doi: 10.1016/j.ceca.2012.11.006. PubMed PMID: 23246155.

42. Verneuil N, Rince A, Sanguinetti M, Posteraro B, Fadda G, Auffray Y, Hartke A, Giard JC. Contribution of a PerR-like regulator to the oxidative-stress response and virulence of Enterococcus faecalis. Microbiology. 2005;151(Pt 12):3997–4004. Epub 2005/12/13. doi: 10.1099/mic.0.28325-0. PubMed PMID: 16339944.

43. Kajfasz JK, Mendoza JE, Gaca AO, Miller JH, Koselny KA, Giambiagi-Demarval M, Wellington M, Abranches J, Lemos JA. The Spx regulator modulates stress responses and virulence in Enterococcus faecalis. Infect Immun. 2012;80(7):2265–75. Epub 2012/04/18. doi: 10.1128/IAI.00026-12. PubMed PMID: 22508863; PMCID: PMC3416481.

44. Muller C, Cacaci M, Sauvageot N, Sanguinetti M, Rattei T, Eder T, Giard JC, Kalinowski J, Hain T, Hartke A. The Intraperitoneal Transcriptome of the Opportunistic Pathogen Enterococcus faecalis in Mice. PLoS One. 2015;10(5):e0126143. Epub 2015/05/16. doi: 10.1371/journal.pone.0126143. PubMed PMID: 25978463; PMCID: PMC4433114.

45. Yang H, Kundra S, Chojnacki M, Liu K, Fuse MA, Abouelhassan Y, Kallifidas D, Zhang P, Huang G, Jin S, Ding Y, Luesch H, Rohde KH, Dunman PM, Lemos JA, Huigens RW, 3rd. A Modular Synthetic Route Involving N-Aryl-2-nitrosoaniline Intermediates Leads to a New Series of 3-Substituted Halogenated Phenazine Antibacterial Agents. J Med Chem. 2021;64(11):7275–95. Epub 2021/04/22. doi: 10.1021/acs.jmedchem.1c00168. PubMed PMID: 33881312; PMCID: PMC8192493.

46. Choby JE, Skaar EP. Heme Synthesis and Acquisition in Bacterial Pathogens. J Mol Biol. 2016;428(17):3408–28. Epub 2016/03/29. doi: 10.1016/j.jmb.2016.03.018. PubMed PMID: 27019298; PMCID: PMC5125930.

47. Ouattara M, Cunha EB, Li X, Huang YS, Dixon D, Eichenbaum Z. Shr of group A streptococcus is a new type of composite NEAT protein involved in sequestering haem from methaemoglobin. Mol Microbiol. 2010;78(3):739–56. Epub 2010/09/03. doi: 10.1111/j.1365-2958.2010.07367.x. PubMed PMID: 20807204; PMCID: PMC2963705.

48. Brugna M, Tasse L, Hederstedt L. In vivo production of catalase containing haem analogues. FEBS J. 2010;277(12):2663–72. Epub 2010/06/18. doi: 10.1111/j.1742-464X.2010.07677.x. PubMed PMID: 20553500.

49. Spirig T, Malmirchegini GR, Zhang J, Robson SA, Sjodt M, Liu M, Krishna Kumar K, Dickson CF, Gell DA, Lei B, Loo JA, Clubb RT. Staphylococcus aureus uses a novel multidomain receptor to break apart human hemoglobin and steal its heme. J Biol Chem. 2013;288(2):1065–78. Epub 2012/11/08. doi: 10.1074/jbc.M112.419119. PubMed PMID: 23132864; PMCID: PMC3542992.

50. Frankenberg L, Brugna M, Hederstedt L. Enterococcus faecalis Heme-Dependent Catalase. Journal of Bacteriology. 2002;184(22):6351–6. doi: 10.1128/jb.184.22.6351-6356.2002.

51. Baureder M, Hederstedt L. Genes important for catalase activity in Enterococcus faecalis. PLoS One. 2012;7(5):e36725. Epub 2012/05/17. doi: 10.1371/journal.pone.0036725. PubMed PMID: 22590595; PMCID: PMC3349705.

52. Winstedt L, Frankenberg L, Hederstedt L, von Wachenfeldt C. Enterococcus faecalis V583 contains a cytochrome bd-type respiratory oxidase. J Bacteriol. 2000;182(13):3863–6. Epub 2000/06/13. doi: 10.1128/JB.182.13.3863-3866.2000. PubMed PMID: 10851008; PMCID: PMC94564.

53. Saillant V, Lipuma D, Ostyn E, Joubert L, Boussac A, Guerin H, Brandelet G, Arnoux P, Lechardeur D. A Novel Enterococcus faecalis Heme Transport Regulator (FhtR) Senses Host Heme To Control Its Intracellular Homeostasis. mBio. 2021;12(1). Epub 2021/02/04. doi: 10.1128/mBio.03392-20. PubMed PMID: 33531389; PMCID: PMC7858072.

54. Coates CJ, Nairn J. Diverse immune functions of hemocyanins. Dev Comp Immunol. 2014;45(1):43–55. Epub 2014/02/04. doi: 10.1016/j.dci.2014.01.021. PubMed PMID: 24486681.

55. Liu M, Huang M, Huang L, Biville F, Zhu D, Wang M, Jia R, Chen S, Zhao X, Yang Q, Wu Y, Zhang S, Huang J, Tian B, Chen X, Liu Y, Zhang L, Yu Y, Pan L, Ur Rehman M, Cheng A. New Perspectives on Galleria mellonella Larvae as a Host Model Using Riemerella anatipestifer as a Proof of Concept. Infect Immun. 2019;87(8). Epub 2019/06/05. doi: 10.1128/IAI.00072-19. PubMed PMID: 31160365; PMCID: PMC6652747.

56. Ge R, Sun X. Iron acquisition and regulation systems in Streptococcus species. Metallomics. 2014;6(5):996–1003. Epub 2014/03/26. doi: 10.1039/c4mt00011k. PubMed PMID: 24663493.

57. Lisiecki P, Wysocki P, Mikucki J. Occurrence of siderophores in enterococci. Zentralbl Bakteriol. 2000;289(8):807–15. Epub 2000/03/08. doi: 10.1016/s0934-8840(00)80006-7. PubMed PMID: 10705612.

58. Sobis-Glinkowska M, Mikucki J, Lisiecki P. [Hydroxamate siderophore effect on growth of enterococci]. Med Dosw Mikrobiol. 2001;53(1):1–7. Epub 2002/01/05. PubMed PMID: 11757399.

59. Lisiecki P, Mikucki J. [Citric acid as a siderophore of enterococci?]. Med Dosw Mikrobiol. 2004;56(1):29–40. Epub 2004/11/05. PubMed PMID: 15524394.

60. Schwyn B, Neilands JB. Universal chemical assay for the detection and determination of siderophores. Anal Biochem. 1987;160(1):47–56. Epub 1987/01/01. doi: 10.1016/0003-2697(87)90612-9. PubMed PMID: 2952030.

61. Keogh D, Lam LN, Doyle LE, Matysik A, Pavagadhi S, Umashankar S, Low PM, Dale JL, Song Y, Ng SP, Boothroyd CB, Dunny GM, Swarup S, Williams RBH, Marsili E, Kline KA. Extracellular Electron Transfer Powers Enterococcus faecalis Biofilm Metabolism. mBio. 2018;9(2). Epub 2018/04/11. doi: 10.1128/mBio.00626-17. PubMed PMID: 29636430; PMCID: PMC5893876.

62. Marcelis JH, den Daas-Slagt HJ, Hoogkamp-Korstanje JA. Iron requirement and chelator production of staphylococci, Streptococcus faecalis and enterobacteriaceae. Antonie Van Leeuwenhoek. 1978;44(3-4):257–67. Epub 1978/01/01. doi: 10.1007/BF00394304. PubMed PMID: 110252.

63. Vebo HC, Snipen L, Nes IF, Brede DA. The transcriptome of the nosocomial pathogen Enterococcus faecalis V583 reveals adaptive responses to growth in blood. PLoS One. 2009;4(11):e7660. Epub 2009/11/06. doi: 10.1371/journal.pone.0007660. PubMed PMID: 19888459; PMCID: PMC2766626.

64. Vebo HC, Solheim M, Snipen L, Nes IF, Brede DA. Comparative genomic analysis of pathogenic and probiotic Enterococcus faecalis isolates, and their transcriptional responses to growth in human urine. PLoS One. 2010;5(8):e12489. Epub 2010/09/09. doi: 10.1371/journal.pone.0012489. PubMed PMID: 20824220; PMCID: PMC2930860.

65. Frank KL, Colomer-Winter C, Grindle SM, Lemos JA, Schlievert PM, Dunny GM. Transcriptome analysis of Enterococcus faecalis during mammalian infection shows cells undergo adaptation and exist in a stringent response state. PLoS One. 2014;9(12):e115839. Epub 2014/12/30. doi: 10.1371/journal.pone.0115839. PubMed PMID: 25545155; PMCID: PMC4278851.

66. Abrantes MC, Kok J, Lopes Mde F. EfaR is a major regulator of Enterococcus faecalis manganese transporters and influences processes involved in host colonization and infection. Infect Immun. 2013;81(3):935–44. Epub 2013/01/09. doi: 10.1128/IAI.06377-11. PubMed PMID: 23297382; PMCID: PMC3584879.

67. Pi H, Helmann JD. Sequential induction of Fur-regulated genes in response to iron limitation in Bacillus subtilis. Proc Natl Acad Sci U S A. 2017;114(48):12785–90. Epub 2017/11/15. doi: 10.1073/pnas.1713008114. PubMed PMID: 29133393; PMCID: PMC5715773.

68. Jakubovics NS. An ion for an iron: streptococcal metal homeostasis under oxidative stress. Biochem J. 2019;476(4):699–703. Epub 2019/03/02. doi: 10.1042/BCJ20190017. PubMed PMID: 30819932.

69. Romero-Saavedra F, Laverde D, Budin-Verneuil A, Muller C, Bernay B, Benachour A, Hartke A, Huebner J. Characterization of Two Metal Binding Lipoproteins as Vaccine Candidates for Enterococcal Infections. PLoS One. 2015;10(8):e0136625. Epub 2015/09/01. doi: 10.1371/journal.pone.0136625. PubMed PMID: 26322633; PMCID: PMC4556446.

70. Obaidullah AJ, Ahmed MH, Kitten T, Kellogg GE. Inhibiting Pneumococcal Surface Antigen A (PsaA) with Small Molecules Discovered through Virtual Screening: Steps toward Validating a Potential Target for Streptococcus pneumoniae. Chem Biodivers. 2018;15(12):e1800234. Epub 2018/09/18. doi: 10.1002/cbdv.201800234. PubMed PMID: 30221472.

71. Lau CK, Krewulak KD, Vogel HJ. Bacterial ferrous iron transport: the Feo system. FEMS Microbiol Rev. 2016;40(2):273–98. Epub 2015/12/20. doi: 10.1093/femsre/fuv049. PubMed PMID: 26684538.

72. Sebulsky MT, Hohnstein D, Hunter MD, Heinrichs DE. Identification and characterization of a membrane permease involved in iron-hydroxamate transport in Staphylococcus aureus. J Bacteriol. 2000;182(16):4394–400. Epub 2000/07/27. doi: 10.1128/JB.182.16.4394-4400.2000. PubMed PMID: 10913070; PMCID: PMC94608.

73. Grigg JC, Ukpabi G, Gaudin CF, Murphy ME. Structural biology of heme binding in the Staphylococcus aureus Isd system. J Inorg Biochem. 2010;104(3):341–8. Epub 2009/10/27. doi: 10.1016/j.jinorgbio.2009.09.012. PubMed PMID: 19853304.

74. Montanez GE, Neely MN, Eichenbaum Z. The streptococcal iron uptake (Siu) transporter is required for iron uptake and virulence in a zebrafish infection model. Microbiology. 2005;151(Pt 11):3749–57. Epub 2005/11/08. doi: 10.1099/mic.0.28075-0. PubMed PMID: 16272396.

75. Morgenthau A, Livingstone M, Adamiak P, Schryvers AB. The role of lactoferrin binding protein B in mediating protection against human lactoferricin. Biochem Cell Biol. 2012;90(3):417–23. Epub 2012/02/16. doi: 10.1139/o11-074. PubMed PMID: 22332888.

76. Bryan-Jones DG, Whittenbury R. Haematin-dependent oxidative phosphorylation in Streptococcus faecalis. J Gen Microbiol. 1969;58(2):247–60. Epub 1969/10/01. doi: 10.1099/00221287-58-2-247. PubMed PMID: 4391229.

77. Holyoake LV, Poole RK, Shepherd M. The CydDC Family of Transporters and Their Roles in Oxidase Assembly and Homeostasis. Adv Microb Physiol. 2015;66:1–53. Epub 2015/07/27. doi: 10.1016/bs.ampbs.2015.04.002. PubMed PMID: 26210105.

78. Pittman MS, Corker H, Wu G, Binet MB, Moir AJ, Poole RK. Cysteine is exported from the Escherichia coli cytoplasm by CydDC, an ATP-binding cassette-type transporter required for cytochrome assembly. J Biol Chem. 2002;277(51):49841–9. Epub 2002/10/24. doi: 10.1074/jbc.M205615200. PubMed PMID: 12393891.

79. Ch’ng JH, Muthu M, Chong KKL, Wong JJ, Tan CAZ, Koh ZJS, Lopez D, Matysik A, Nair ZJ, Barkham T, Wang Y, Kline KA. Heme cross-feeding can augment Staphylococcus aureus and Enterococcus faecalis dual species biofilms. ISME J. 2022. Epub 2022/05/20. doi: 10.1038/s41396-022-01248-1. PubMed PMID: 35589966.

80. Kajfasz JK, Katrak C, Ganguly T, Vargas J, Wright L, Peters ZT, Spatafora GA, Abranches J, Lemos JA. Manganese Uptake, Mediated by SloABC and MntH, Is Essential for the Fitness of Streptococcus mutans. mSphere. 2020;5(1). Epub 2020/01/10. doi: 10.1128/mSphere.00764-19. PubMed PMID: 31915219; PMCID: PMC6952196.

81. Kristich CJ, Manias DA, Dunny GM. Development of a method for markerless genetic exchange in Enterococcus faecalis and its use in construction of a srtA mutant. Appl Environ Microbiol. 2005;71(10):5837–49. Epub 2005/10/06. doi: 10.1128/AEM.71.10.5837-5849.2005. PubMed PMID: 16204495; PMCID: PMC1265997.

